# Small extracellular vesicles secreted by Candida albicans hyphae have highly diverse protein cargoes that include virulence factors and stimulate macrophages

**DOI:** 10.1101/2020.10.02.323774

**Authors:** Raquel Martínez-López, Maria Luisa Hernáez, Esther Redondo, Guillermo Calvo, Sonja Radau, Concha Gil, Lucía Monteoliva

## Abstract

Extracellular vesicles (EVs) have been described as mediators of microorganism survival and interaction with the host. In *Candida albicans*, a relevant commensal fungal pathogen, the dimorphic transition is an important virulence trait in candidiasis. We have analyzed EVs secreted by yeast (YEVs) or hyphal cells (HEVs) from *C. albicans*, finding interesting differences in both size distribution and protein loading. In general, HEVs were smaller and carried a much more diverse protein cargo than YEVs, including most of the proteins identified in YEVs, which were mainly cell surface proteins. Virulence factors such as phospholipases, aspartic proteases (Saps), adhesins and invasins, and the precursor protein of candidalysin toxin Ece1p were also detected. HEVs were also enriched in proteasomal and ribosomal proteins, and in enzymes from amino acid biosynthetic pathways, all involved in protein metabolism, as well as proteins related to intracellular protein transport and components of the ESCRT pathway related to exosome biogenesis. Both types of EV presented immune reactivity with human sera from patients suffering invasive candidiasis. In our conditions, only HEVs could elicit the release of TNFα by activated macrophages. This first analysis of *C. albicans* HEVs shows their relevance to pathogenesis and possible new diagnostics or treatments.

## INTRODUCTION

*C. albicans* can be found as a commensal fungus of humans, mainly on skin and mucosal surfaces such as the oral cavity, gastrointestinal tract, and vagina. However, when host immunity is disrupted, *C. albicans* can cause an infection known as candidiasis, which can go from superficial candidiasis to life-threatening invasive candidiasis.^1, 2^ Cancer therapies and organ transplantation, which have increased exponentially in recent years, are some of the conditions associated with immunosuppression that favors invasive candidiasis. The *C. albicans* yeast-to-hypha transition is highly studied, since it is critical for virulence. The hyphal morphology is generally considered to be more related to the invasion of host tissues, while the yeast morphology is more suited to bloodstream dissemination or surface commensalism.^3^ Proteomic studies of *C. albicans* dimorphism have used a variety of approaches, ranging from analyses of cytoplasmic and cell-wall proteins from yeast cells, hyphae, and biofilms to quantitative analysis of the proteome during the yeast-to-hypha transition.^4, 5^ Novel strategies have also been developed, such as the one described by Hernaez et al. based on “cell shaving” of live *C. albicans* cells. This was applied to both yeast and hyphae and led to interesting findings such as the identification of novel proteins involved in cell wall integrity, the yeast-to-hypha transition, and stress response and/or host-pathogen interaction.^6, 7^ Moreover, a similar strategy was used to decipher not only *C. albicans* proteins but also human serum proteins that were linked to the hyphal surface when yeast cells of *C. albicans* were incubated with serum promoting their switch to hyphae.^8^ Several proteins classically considered cytoplasmic because they lack signal peptides, including components of metabolic pathways, chaperones, and ribosomal proteins, have long been identified in proteomic studies as residing in the *C. albicans* cell wall or as part of the *C. albicans* secretome.^9-13^ Some secretory pathways that are alternatives to the endoplasmic reticulum-Golgi pathway for signal-peptide-containing proteins have started to emerge.^10, 11, 14^ In addition, the existence of extracellular vesicles (EVs) in gram positive and gram negative bacteria, and in fungi is being recognized.^15^ Nowadays it is widely accepted that cells from almost every type of organism secrete these nano- to micrometer-scale lipid-bilayer-delimited vesicles.^15^ In *C. albicans*, Anderson et al. demonstrated the existence of vesicle-like compartments in cell wall pimples from opaque cultures of *C. albicans* cells in 1990.^16^ *C. albicans* EV were first isolated and observed by transmission electron microscopy (TEM) in 2008 by Albuquerque et al., who demonstrated the presence of bilayered compartments similar to those initially described for *C. neoformans* and *H. capsulatum*.^17, 18^ EVs of *C. albicans* yeast cells were later further analyzed to unravel their composition and implications for human immune responses in wild-type and mutant strains.^9, 19-21^ All this work has been extensively reviewed by Gil-Bona et al.^10^

Human EVs are well-studied and have been classified as apoptotic bodies, ectosomes, or exosomes, depending upon their cellular origin and size.^22^ Apoptotic bodies are the largest (50– 5000 nm in size) and are derived from apoptotic cells. Ectosomes, also called microvesicles, are generated by outward budding from the plasma membrane, followed by pinching off and release to the extracellular space, resulting in EVs ranging from 100 to 1000 nm in size.

Exosomes are the smallest EVs (30 to 150 nm), and these structures originate from endosomal compartments.^22-24^ EVs shuttle bioactive molecules involved in many processes including cell-cell communication, host-pathogen interactions, and even the sharing of microbial community resources, in the case of microbial EVs. For example, *Cryptococcus neoformans* extracellular vesicles contain its major virulence factor, the capsular polysaccharide glucuronoxylomannan.^25^ It has been reported that EVs secreted by wild type *C. albicans* biofilm are able to rescue the antifungal resistance of a defective biofilm produced by cells carrying mutations in genes encoding orthologues of endosomal sorting complexes required for transport (ESCRT) subunits.^26^ Other authors have proposed the use of these EVs as therapeutic carriers of drugs and metabolites, since the internalization of EVs secreted by different microorganisms and different mammalian cells by both microorganisms and mammalian cells of different host tissues has been widely proven in several recent research papers.^27-29^

In this context, and given the established contribution of EVs to key physiological aspects of cells from all kingdoms, we isolated and characterized EVs secreted by *C. albicans* of both major morphologies, yeast and hyphae, to better understand the mechanisms underlying the enhanced virulence associated with the morphologic transition. Differences in EV size and physical properties were analyzed by means of transmission electronic microscopy (TEM) and dynamic light scattering (DLS). Protein cargoes were analyzed using LC-MS. The more interesting differences were observed in the proteomic analysis, suggesting that hyphal EVs (HEVs) differ in their biogenesis and function from yeast EVs (YEVs).

## EXPERIMENTAL SECTION

### Microorganisms and culture conditions

The *C. albicans* clinical isolate SC5314 ^30^ was used in this work. It was grown on YPD agar plates (1% D-glucose, 1% Difco Yeast Extract, and 2% agar) overnight at 30 °C prior to the experiment. Two isolated colonies were used to inoculate 200 mL of liquid SD medium (20 g/L de glucose, 5 g/L ammonium sulphate, 1.7 g/L yeast nitrogen base, 1.92 g/L synthetic amimoacid mixture minus uracile (Formomedium) supplemented with 0.1 g/L uracil). *C. albicans* culture was growing during 6 hs at 30°C and 180 rpm. Cells were then collected by 10 min centrifugation at 2 500 rpm in an Eppendorf 5810R centrifuge, supernatant discarded and cells washed with 1 ml of Phosphate saline buffer (PBS) and collected again by 3 min centrifugation at 5 000 rpm in a microcentrifuge Heraeus Fresco 21™ (Thermo Scientific). Cells were then counted on a Neubauer chamber to inoculate 10^6^ cells in 1 L of the different media used to obtain the specific morphology.

For yeast morphology 1 L of YNBS (5 g/L ammonium sulphate, 1.7 g/L yeast nitrogen base, 20 g/L sucrose) was supplemented with 75 mM of tartaric acid adjusted to pH 4.

For hyphal morphology 1 L of YNBS was supplemented with 75 mM of 3-(N-morfolin) propanosulfónic acid (MOPS) adjusted at pH 7,4 and 5 mM de N-acetylglucosamine (N-AcGlc).^31, 32^

### *C. albicans* cell viability measurement

Prior to EVs isolation, 1mL of yeast and hyphae culture media was treated with propidium iodide (PI) to test cell viability. PI is non-permeable to intact cell membranes but can enter death cells with compromised membranes dying DNA molecules. 10^6^ cells were incubated with 10 µl of PI 5 mM (Fluka) and the positive fluorescence cells were counted under a fluorescence microscope at λ=450 nm. 70% ethanol-PBS treated cells were used as positive death cells control. At least 500 cells of each sample were counted to calculate the % of inviable PI stained cells.

### Isolation of Extracellular Vesicles

Three independent experiments were done with yeast and hyphal cultures. The isolation of EVs was done according to Gil Bona.^19^ All the process was conducted at 4°C. In brief, the supernatants from 1 L of yeast and hyphal specific culture media grown during 16h at 37°C and 180 rpm, were collected by 20 min centrifugation at 8 000 rpm at 4°C in a Beckman Coulter J2-HS centrifuge using the JA-10 rotor. Supernatants were then filtrated using a 0,45 µm filter to ensure the elimination of all the cells and cell debris. One protease-inhibitor tablet (Pierce™ EDTA-free, Thermo Fisher) along with 1mL of Phenylmethanesulfonyl fluoride (PMSF) was added to each of the 1L filtrated supernatants. These supernatants were concentrated afterwards using a Centricon Plus-70 filter (cutoff filter 100 kDa, Millipore) by centrifugation at 2500 rpm in an Eppendorf 5810R centrifuge to a final volume of 8 mL. The concentrated supernatants were subsequently ultra-centrifuged at 100 000 g (34200 rpm) for 1 h at 4 °C in a Beckman Optima XL-90 using 90 Ti rotor. The pellets containing the isolated EVs were washed twice with PBS and solubilized in 50 µl of triethylammonium bicarbonate (TEAB) buffer 0.5 M. Protein concentration was measured using Bradford reactive (BioRad) following manufacture instructions.

### Transmission Electron Microscopy (TEM)

Electron microscopy (TEM) was used to visualize isolated EVs from both yeast and hyphal cell morphologies. Samples were fixing for 2 h at room temperature in a buffer containing 2.5% glutaraldehyde and 0.1 M cacodylate and then were incubated overnight at 4 °C in 4% paraformaldehyde, 1% glutaraldehyde, and 0.1% PBS. After that, the samples were treated with 2% osmium tetroxide (TAAB Laboratories, U.K.) for 90 min, serially dehydrated in ethanol, and embedded in EMBed-812 resin (Electron Microscopy Sciences). Thin sections (50−70 nm) were obtained by ultra-cut and observed in a JEOL JEM 1010 transmission electron microscope operating at 100 kV. Pictures were taken with Megaview II camera. TEM images were analyzed with Soft Imaging Viewer Software. TEM was carried out in Electronic microscopy facility of the Complutense University of Madrid (ICTS)-UCM.

### Analysis of vesicles by Dynamic light scattering (DLS)

EV sizes (z-average diameter) were measured by dynamic light scattering (DLS) using a ZetaSizer (Nano ZS, Malvern). Vesicles in a liquid phase undergo Brownian motion, and this produces light-scattering fluctuations, which give information on the size and heterogeneity of a sample. Three biological replicates of EVs (from both type of cells, yeast, and hyphae) were transferred to a disposable cuvette, and 10 measurements for each were performed with refractive index at 1.33 and absorption at 0.01. Data analysis was performed using the Zetasizer Software 7.11 (Malvern). DLS was carried out at the spectroscopy and correlation facility of the Complutense University of Madrid (UCM).

### Whole cell lysates (WCL) protein extraction

Three independent experiments were done with yeast and hyphal cultures. All the processes were conducted at 4°C. 20 mL of yeast and hyphal specific culture media grown during 16 hs at 37°C and 180 rpm were collected in a 50 mL falcon by 10 min centrifugation at 2 500 rpm at 4°C in an Eppendorf 5810R centrifuge. The cell pellets were subsequently washed twice with 20mL of ice-cold PBS and transferred to a 2 mL Eppendorf. Equal volumes of 0.45-mm glass beads were added. The cell pellets were disrupted in 500 µL of lysis buffer (50 mM Tris-HCl pH7.5, 1 mM EDTA, 150 mM NaCl, 1 mM DTT) with 1 mM PMSF and protease inhibitor cocktail tablets (Roche) by vigorous shaking in a fast prep cell breaker (Bio 101; level of 5.5, 5 times for 30). Cell debris and glass beads were removed by centrifugation (13 000 rpm for 15 min) and the cell extracts were collected in a new Eppendorf tube. Protein quantification was performed using the Bradford assay (Bio-Rad, Hercules, CA, USA) and protein samples were stored at −80 °C.

### SDS-PAGE and Western Blotting

EVs protein extracts (30 μg of each) were denatured by heating for 5 min at 99 °C in SDS containing buffer (4% SDS, 100 mM Tris HCl pH 6.8, 20% glycerol, 0.2% bromophenol blue and 20% DTT). Protein samples were separate in 10% SDS-polyacrylamide gel using the Miniprotean II electrophoresis system (Bio-Rad). The gel was stained with a fixative solution of 40% MeOH, 10% acetic acid v/v and brilliant Coomassie blue (BioRad G250). For Western blotting, 30 μg of EVs protein extracts were separated in 10% SDS-polyacrylamide gels, transferred to nitrocellulose membranes, and blocked in 5% milk PBS. Western blots were probed with sera from patients suffering invasive candidiasis at a dilution of 1:3000. After an overnight incubation with the sera, membranes were washed five times with 0.1% Tween-20 containing PBS, and then incubated with fluorescently labeled secondary antibodies at a dilution of 1/1000 (IR Dye 800 goat antihuman IgG (LI-COR Biosciences). The Western blotting was performed with the Odyssey system (LI-COR Biosciences, Nebraska, USA).

### Digestion and desalting of peptides

In gel protein digestion is useful to eliminate contaminants that could interfere with MS/MS analyses. For this, twenty-five micrograms of each protein extract were concentrated in a stacking gel, and protein bands were cut from the acrylamide gel for in gel trypsin digestion. Briefly, cut protein bands were first DTT reduced (Sigma-Aldrich, St. Louis, MO, USA) to be subsequently treated with Iodacetamide for protein alkylation (Sigma-Aldrich, St. Louis, MO, USA) and ultimately digested with 1.25 µg of recombinant Trypsin (sequencing grade; Roche, Mannheim, Germany) overnight at 37 °C (Sechi and Chait, 1998). A C18 reverse phase chromatography was used for desalting and concentration of the peptides from the digested proteins (POROS R2, Applied Bioystems), being afterwards eluted with 80% acetonitrile/0.1% trifluoroacetic acid (Thermo Fisher Scientific). Elution buffer was then evaporated in Speed-vac (Thermo Fisher Scientific, Rockford, IL, USA), and the freeze-dried samples resuspended in 2% Acetonitrile, 0.1% formic acid (Thermo Fisher Scientific, Rockford, IL, USA) before the Nano liquid chromatography coupled with mass spectrometry in tandem (LC–MS/MS) analysis.

### LC–MS/MS

The desalted peptides were analyzed by a reversed phase liquid chromatography electrospray ionisation tandem mass spectrometry (RP-LC-ESI-MS/MS) in an Ultimate 3000 nLC (Thermo Fisher Scientific) coupled to the Orbitrap Fusion® Lumos® Tribrid® (Thermo Fisher) (an ultrahigh-field mass orbitrap analyzer) through the Nano-Easy spray source (all from Thermo Scientific, Bremen, Germany). Peptides were loaded first onto an Acclaim PepMap 100 Trapping column (Thermo Scientific, 20 mm 75 µm inner diameter (ID), 3 µm of C18 resin with 100 Å pore size, Thermo Scientific, Germering, Germany) using buffer A (mobile phase A: 2% acetonitrile, 0.1% formic acid) and then separated and eluted on a C18 resin analytical column NTCC (50 cm 75 µm ID, 3 µm C18 resin with 100 Å pore size, Nikkyo Technos Co., Ltd., Tokyo, Japan) with an integrated spray tip. A 95 min gradient of 5–27% Buffer B (100% acetonitrile, 0.1% formic acid), from 27% to 44% in 5 min and finally 10 min more until 95% in Buffer A at a constant flow rate of 0,3 µl/min.

All data were acquired using data-dependent acquisition (DDA) and in positive mode with Xcalibur 4.0 software (Thermo Fisher Scientific Inc., USA). For the MS2 scan, the top 15 most abundant precursors with charges of 2–7+ selected in MS 1 scans were selected for higher energy collisional dissociation (HCD) fragmentation with a dynamic exclusion of 60 s. The MS1 scans were acquired at a m/z range of 375–1500 Da with a orbitrap mass resolution of 120000 and an automatic gain control (AGC) target of 4E5 at a maximum Ion Time (ITmax) of 50 ms. The threshold to trigger MS2 scans was 5E3; the normalized collision energy (NCE) was 30%; the resolved fragments were scanned at a mass resolution of 30,000 and an AGC target value of 1E4 in an ITmax of 60 ms.

### Protein identification

Peptide identifications from raw data were carried out using the Mascot v. 2.6.1 (MatrixScience, London, UK) search engine through the Protein Discoverer 2.4 Software (Thermo Fisher Scientific, Waltham, MA, USA). A database search was performed against Candida albicans CGD21 (6209 sequences) from http://www.candidagenome.org. The following parameters were used for the searches: tryptic cleavage, up to two missed cleavage sites allowed, and tolerances of 10 ppm. for precursor ions and 0.02 Da for MS/MS fragment ions, and the searches were performed allowing optional methionine oxidation and acetyl protein N-terminal and fixed carbamidomethylation of cysteine. A search against the decoy database (integrated decoy approach) was used to calculate the false discovery rate (FDR). The Mascot Scores were adjusted by a percolator algorithm. The acceptance criteria for protein identification were an FDR <0.01. The mass spectrometry proteomics data have been deposited to the ProteomeXchange Consortium via the PRIDE [1] partner repository with the dataset identifier PXD021488 and PXD021504. As an estimation of the relative protein abundances the normalized spectral abundance factor (NSAF) was used, and the average of the normalized values was calculated.^33^

### Bioinformatic Analysis

We used the Candida genome database (CGD, www.candidagenome.org) for the analyses. Proteins that were identified in at least two replicates with at least two peptides in one of them were used for the analysis. Venn diagrams were prepared using the Venn tool available in the program FunRich 3.1.3.^34^ The Go enrichment analysis of the set of proteins identified in yeast or hyphal EVs were done using Gene Ontology (GO) annotation application (http://www.candidagenome.org/cgi-bin/GO/goTermFinder) from CGD and the GO enrichment analysis from the FunRich 3.1.3 program which is based on a Uniprot database and uses protein homologues from all fungi.^34^

Metabolic pathways were retrieved from KEEG database (https://www.genome.jp/kegg/).^35^

### THP-1 Cell Culture and Macrophage Differentiation

THP-1 cells (human acute monocytic leukemia cell line) were grown and maintained in Dulbecco’s modified eagle’s medium (DMEM) supplemented with antibiotics (penicillin 10000 U/mL− streptomycin 10000 U/mL), 2 mM L-glutamine, and 10% heat-inactivated fetal bovine serum (FBS). THP-1 cultures were incubated in a humidified atmosphere containing 5% CO_2_ at 37 °C. 24-well plastic plates were seeded with THP-1 cells at a density of 30 ⨯ 10^4^ cells per well in complete medium after being treated with 30 ng/mL phorbol 12-myristate 13-acetate (PMA; Sigma-Aldrich). These 24-well plastic plates where then incubated for 48 hs to induce maturation mediated by PMA toward adherent macrophage-like cells. After this 48 hs period, the medium containing PMA was replaced with fresh medium without PMA to remove unattached cells.

### Determination of cytokine production

For cytokines measurements, differentiated macrophages from the THP1 cell line were incubated for 8 and 24 hs with or without 5µg of YEVs or HEVs. As a positive control, macrophages were treated with lipopolysaccharide (LPS; 100 ng/mL). After the corresponding incubation period, supernatants from THP-1 macrophages (untreated, LPS (1000 ng/ml), and YEVS or HEVS treated) were collected. They were tested for cytokine production by ELISA using matched paired antibodies specific for IL-12p40, TNF-α and IL-10 (Immunotools), and according to the manufacturer’s instructions. Cytokine concentration was measured spectrophotometrically at 450 nm in a total of 3 biological replicates.

### Macrophage damage assay

A colorimetric assay based on the measurement of LDH activity released by damaged cells was used (Roche). Experiments were performed in 96-well plate and the manufacturer’s instructions were followed. Briefly, Lysis buffer was added to the positive control cells 15 min before the end of the incubation time. To determine LDH activity, 100 µl of reaction mixture, (catalyst and dye solution) was added to each well on the 96-well plate and incubated (protected from the light) for up to 30 min at room temperature. After this, 50 µl of stop solution (H_2_SO_4_ 2M) was added to each well and absorbance at 490 nm was measured for each one. Cytotoxicity was calculated as follows:

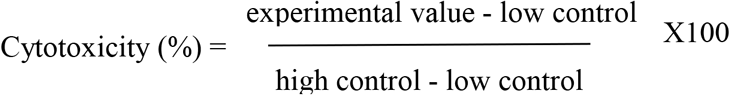

Three biological replicates were performed.

## RESULTS

We have accomplished the study of YEVs and HEVs secreted by *C. albicans* cells. The use of YNBS cultures in which the pH was varied by addition of either tartaric acid (pH 4) or a combination of MOPS and N-acetyl glucosamine (pH 7) allowed the culture of pure yeast and hyphae morphologies respectively. Acidic pH favors yeast morphology while neutral pH together with MOPS and N-acetyl glucosamine ensures the morphological switch to the filamentous hyphal form (Figure S1). EVs were obtained from culture media supernatants after a 16-h incubation period. Prior to EV isolation, morphological purity was assessed by microscopy and the cultures were observed to be around 99% pure yeast or hyphae. In addition, a viability assay was conducted using propidium iodide (PI) staining to check for the absence of cells with altered permeability, as their released intracellular contents could ultimately alter the protein content of the supernatant. All samples showed a PI staining rate below 1% (Figure S1). *C. albicans* cultures were centrifuged at 8000 rpm and subsequently filtrated using a 0.45-µm filter to remove the yeast or hyphal cells. The 1 L volume was concentrated to 8 mL with an Amicon filter with a 100-kD cut-off. Vesicles were collected after a 1-h ultracentrifugation (100 000 × *g*).

### 1. Extracellular vesicles secreted by yeast and hyphae differ in size

YEVS and HEVs collected from culture supernatants were analyzed using DLS and TEM (Figure1a and 1b). Based on the size-distribution by intensity pattern obtained for the EVs, we observed that the majority of YEVs collected were significantly bigger than the HEVs, most of them being in the range of 400-500 nm. Nevertheless, there was also a small percentage of YEVs similar to most of the HEVs in size, with a peak at 100 nm. HEVs presented a less homogeneous population, with a size distribution ranging from 50-450 nm, with the higher proportion of HEVs in the range of 100-200 nm. This difference in size can also be appreciated in the TEM analysis (Figure1a and 1b) where the generally bigger size of YEVs was clear.

**Figure 1.**
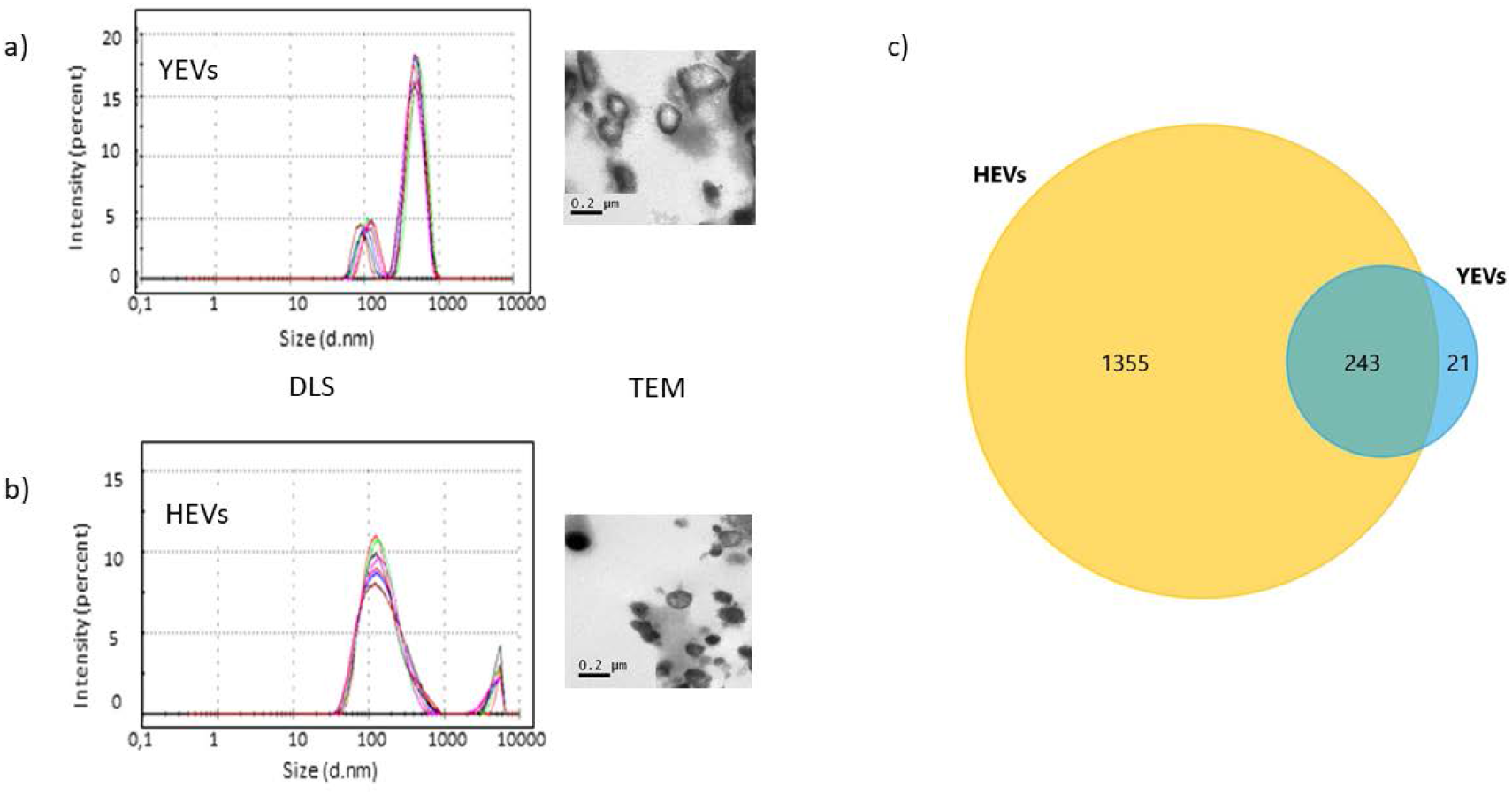
Size –distribution by intensity pattern (DLS) and appearance (TEM) of a) YEVs and b) HEVs. c) Venn diagram showing number of identified proteins which are common or exclusive to EVs from each cell morphology.

### 2. HEVs carry a much more highly diverse protein cargo than YEVs

The protein cargoes of EVs from cells of each morphology were identified by means of LC-MS/MS proteomic analysis. A great difference in the number of identified proteins depending on the EVs origin was immediately conspicuous. Up to 2034 different proteins were identified from HEVs, counting all the different proteins from the three biological replicates carried out, while only 349 proteins were identified in the three biological replicates from YEVs. In terms of biological significance, we took into consideration only proteins that were identified in at least two biological replicates with at least two peptides in one of the replicates and a q-value < 0.01 (Table S1). With these criteria, the number of proteins in HEVs was 1598, and in YEVs was 264, and the significant difference in protein diversity displayed by the two types of EVs was retained. Of the 264 proteins identified in YEVs, 243 were also identified in HEVs (Figure 1c). To carry out the comparative analysis we established 3 sets of proteins: proteins exclusively identified in YEVs, proteins identified in both types of vesicle, and proteins identified exclusively in HEVs (Figure 1c).

Identified proteins were listed according to their normalized spectral abundance factor (NSAF) values to reveal their relative abundance in EVs from each cell morphology. It is worth mentioning that the NSAF value is normalized by considering all the proteins identified in each sample, so that in general, proteins in samples with a higher number of identified proteins will have smaller NSAF values (Figure 2). The comparative analysis of these data is commented below.

**Figure 2.**
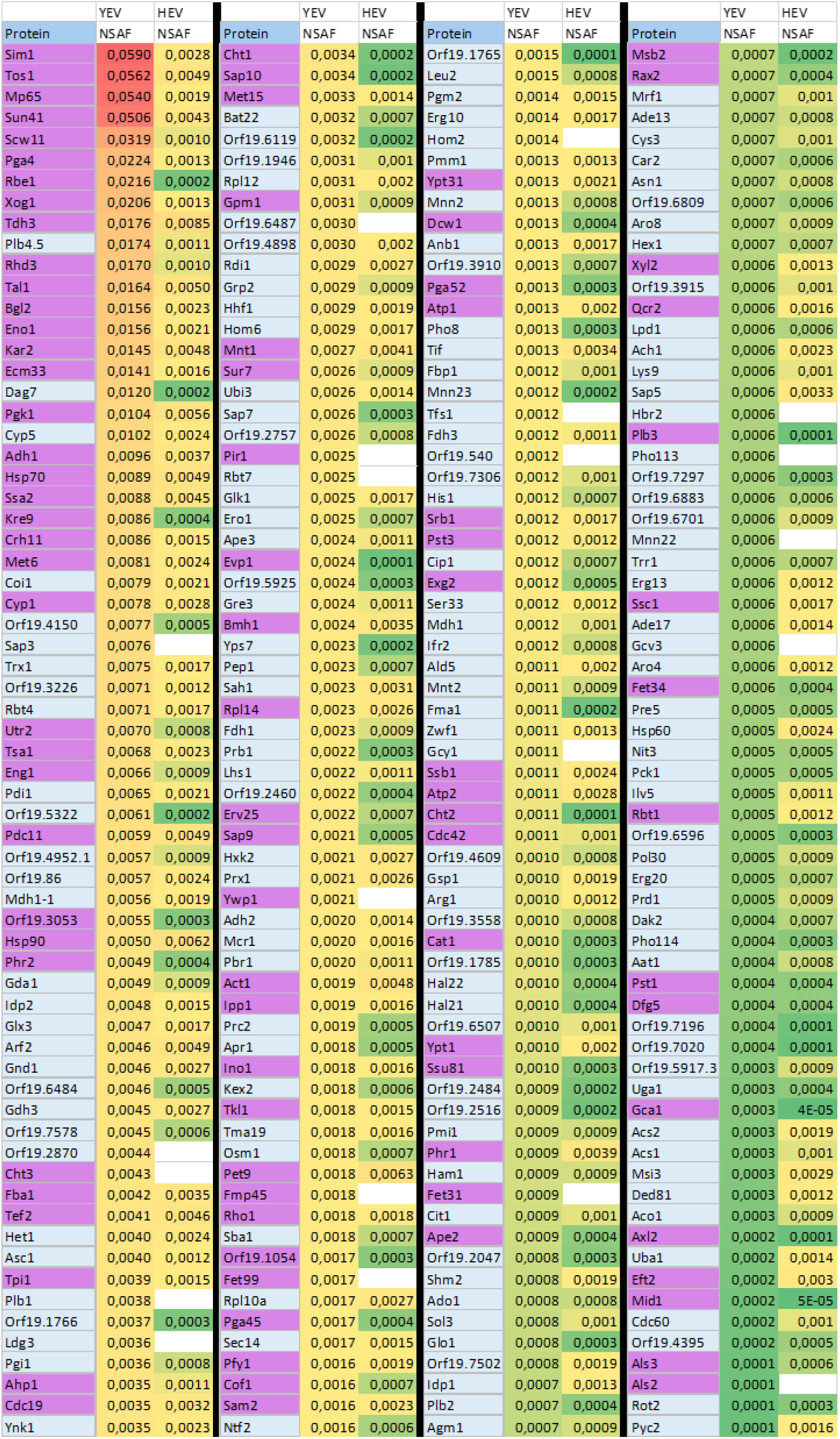
List of YEVs proteins according to decreasing NSAF values and compared with their relative abundance in HEVs. Proteins exclusively identified in YEVs show a blank in the HEV column. Conditional coloring is applied according relative abundance (red the most abundant and green the least). Proteins associated to the cell periphery according to the CGD are shadowed in purple.

#### 2.1 Both YEVs and HEVs are enriched in proteins associated with the cell periphery and membranes

As mentioned above, most of the proteins identified in YEVs were also identified in HEVs. In Figure 2, YEVs proteins are listed according to decreasing NSAF or relative abundance value and compared with their relative abundance in HEVs. In general, proteins common to HEVs and YEVs had similar relative NSAF values in the two types of EV, with the exception of some such as Rbe1, Dag7, Phr2, Kre9, Cht1, Sap10, Sap7, Bat22, Yps7, Pep1or Prb1, and some orfs (Orf19.5322, Orf19.3053, Orf19.6484, Orf19.1766, Orf19.6119, Orf19.6741, Orf19.5925) that were among the 100 most abundant proteins in YEVs but had low NSAFs in HEVs (Figure 2).

GO enrichment analysis from the Candida Genome Database (CGD) of proteins common to both type of EVs clearly revealed, in the cell component category, a high degree of enrichment in proteins from extracellular regions (including the cell surface, cell wall, or biofilm matrix) or anchored to the plasma membrane. FunRich analysis, which is based on a Uniprot database and uses protein homologues from all fungi, was also carried out (Figure 3). As can be observed in Figure 3a, the FunRich categorization of component enrichment was in accord with the CGD analysis.

**Figure 3.**
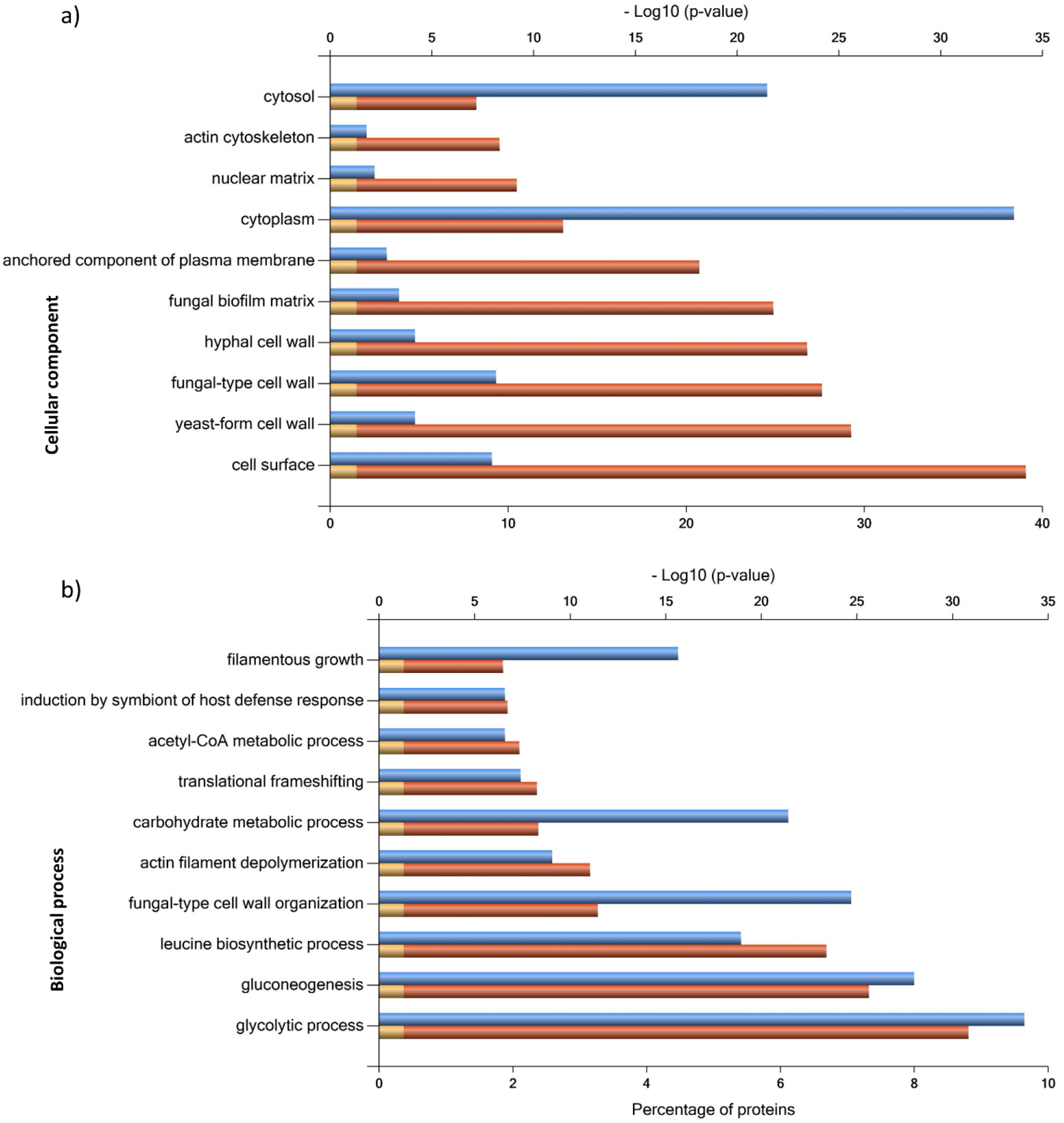
FunRich categorization of component (a) and biological process (b) enrichment of proteins identified in both, YEVs and HEVs. p-value for significance is <0,001.

These proteins included typical cell wall proteins already described in numerous works as well as cell surface-associated proteins such as glycolytic enzymes (Eno1, Tdh3, Pgk1) (Figure 2).

On the other hand, the GO enrichment analysis of the 38 proteins exclusive to or more abundant in YEVs (also identified in HEVs but with low NSAF values) listed in Table S2 and the 1355 proteins exclusively identified in HEVs showed that regarding the first ones, the enrichment in proteins of some extracellular cell components (cell surface or cell wall proteins) was similar to what was observed with proteins common to both types of EV (Figure S2). In contrast, proteins exclusively identified in HEVs showed a completely different enrichment pattern in both the cellular component and biological process categories (Figure 4), as described below.

**Figure 4.**
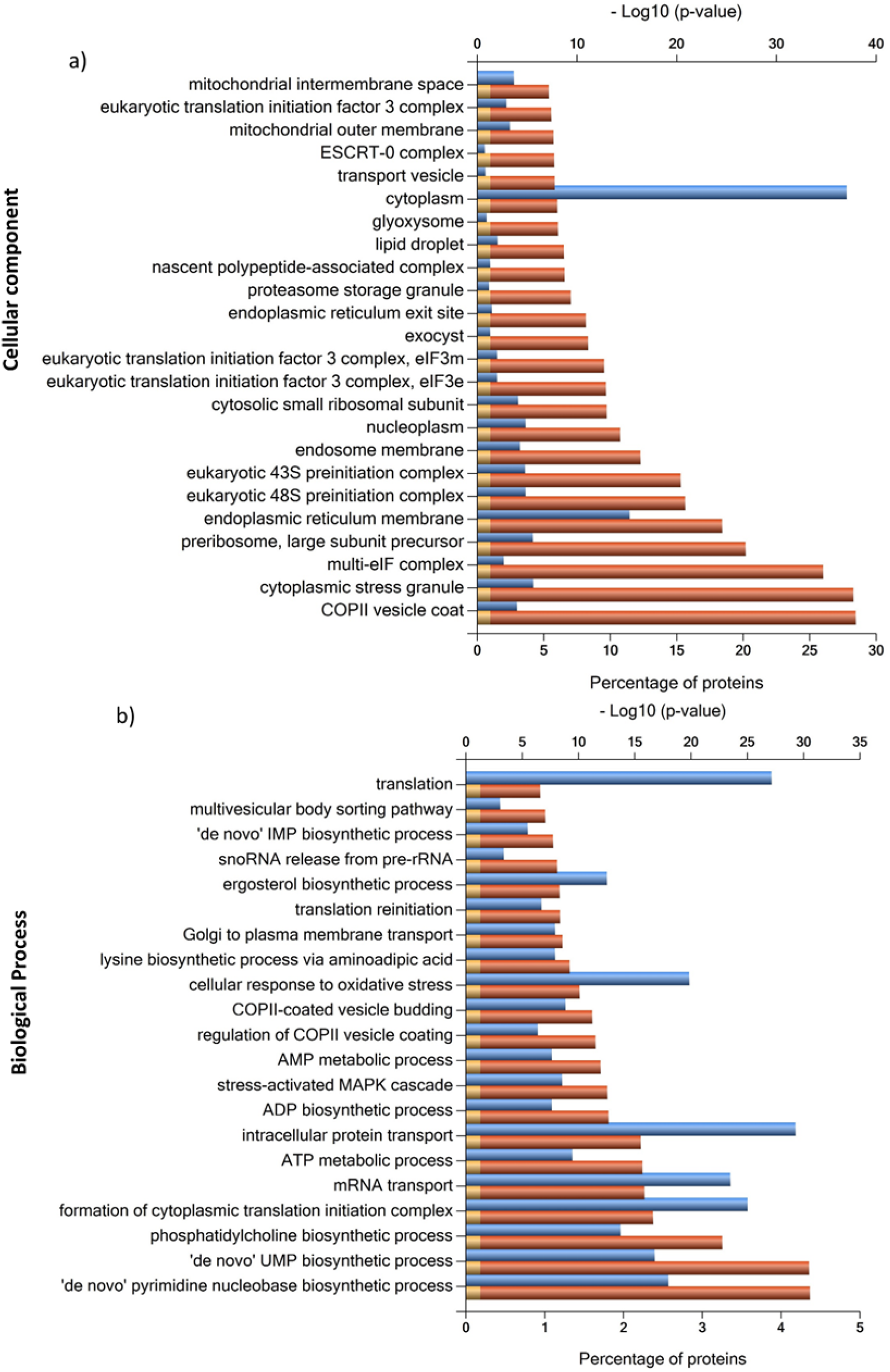
FunRich categorization of component a) and biological process b) enrichment of proteins exclusively identified in HEVs. p-value for significance is <0,001.

#### 2.2 HEVs are enriched in proteins involved in protein metabolism, amino acids synthesis, and *de novo* purine and pyrimidine biosynthesis

GO enrichment analysis of cellular component of proteins exclusively identified in HEVs revealed a predominantly cytoplasmic association, in clear contrast to the enrichment in extracellular component proteins identified in both EVs or exclusively identified or enriched in YEVs (Figure 3 y 4).

Also, in proteins identified only in HEVs there was remarkably high enrichment in GO categories related to protein metabolism, due to the presence of many proteins belonging to multi-eIF, 43S preinitiation, ribosome, and proteasome complexes (Figure 4a). Regarding proteasome structure, we could identify 16 (80%) of the 20 proteins that form the regulatory particle of the proteasome and 12 (92%) of the14 proteins that form the 20S particle core of the proteasome (Figure S3). Furthermore, five of nine proteins described as being associated to the proteasome, such as Ubp6 (a deubiquitinating enzyme), Ubc4, Ecm29, Rad23, and Ddi1, were also identified as exclusively HEVs protein cargo.

We identified 49 ribosomal proteins, including 26 (79%) of the 33 proteins that forms the cytosolic small subunit of the ribosome. The presence in HEVs of these proteins of the cytosolic small subunit of the ribosome together with most of the proteins from the multi-eIF complex lead to the identification of almost the entire 48S preinitiation complex. This complex is involved in translational initiation, which makes translation one of the biological processes by which HEVs are significantly enriched (Figure 4b).

Special attention should be given to the high number of proteins involved in the synthesis of different amino acids (aa) within the protein cargo of HEVs. In fact, the HEVs protein cargo included all the enzymes necessary for the synthesis of valine, leucine, isoleucine, cysteine, serine, glycine, methionine, threonine, alanine, proline, lysine, tyrosine, and glutamine from fructose-6P. We also identified all but one or two of the enzymes necessary for the biosynthesis of histidine, tryptophan, phenylalanine, and asparagine in HEVs.

Also worth mentioning is the high significance of purine and pyrimidine biosynthetic processes among the biological processes represented by proteins exclusively identified in HEVs (Figure 4b). All the purine metabolic enzymes involved in the route from phosphoribosyl pyrophosphate (PRPP) to inosine monophosphate (IMP) and then to adenosine monophosphate (AMP) were identified in HEVs, while only three of them (Ade13, Ade17, Ade6) were identified in YEVs.

All the pyrimidine metabolic enzymes involved in the route from L-glutamine to uridine monophosphate (UMP) were exclusively identified in HEVs.

#### 2.3 HEVs are significantly enriched in vesicular transport-related proteins

It is also remarkable that HEVs are highly enriched in proteins exclusively identified in them that belong to cellular components of the classical protein secretion pathway, such as ER, Golgi or COPII vesicles coats, or other components involved in vesicular traffic, such as components of the the ESCRT-0 complex, the endosome membrane, or the exocyst (Figure 4a). Thus, many processes related to protein and vesicle transport had significant p-values only in the HEVs protein set (Figure 4b) and were completely absent or with nonsignificant p-values in YEVS (Figure S2). In this sense, HEVs were enriched in class e-vacuolar protein sorting (Vps) and other selected proteins required for multivesicular body (MVB) sorting cargo, and proteins related to ER-Golgi vesicle transport and exocytosis, in addition to COPI and COPII vesicle-related proteins (Table 1).

**Table 1.** List of proteins identified in HEV and YEVs and related to pathways and structures involved in vesicular transport (according to the CGD).

| Protein processing in endoplasmic reticulum: map04141 |  |  |  |
| --- | --- | --- | --- |
| Protein | Description according CGD | HEVs | YEVs |
| Cdc48 | Putative microsomal ATPase; plasma membrane-localized | ✓ | ✗ |
| Cwh41 | Processing alpha glucosidase I, involved in N-linked protein glycosylation and assembly of cell wall beta 1,6 glucan | ✓ | ✗ |
| Emp46 | Protein similar to *S. cerevisiae* Emp46, an integral membrane component of ER-derived COPII-coated vesicles; functions in ER to Golgi transport | ✓ | ✗ |
| Ire1 | Putative protein kinase; role in cell wall regulation | ✓ | ✗ |
| Jem1 | Functional homolog of *S. cerevisiae* Jem1p, which acts with Scj1p and Kar2p (BiP) in protein folding and ER-associated degradation of misfolded proteins | ✓ | ✗ |
| Kar2 | Similar to Hsp70 family chaperones; role in translocation of proteins into the ER | ✓ | ✓ |
| Lhs1 | Protein similar to *S. cerevisiae* Hsp70p; predicted Kex2p substrate | ✓ | ✓ |
| Mns1 | member of ER localized glycosyl hydrolase family 47; ER form is converted by Kex2 to cytosolic form | ✓ | ✗ |
| Npl4 | Putative ubiquitin-binding protein; regulated by Gcn2p and Gcn4p | ✓ | ✗ |
| Ost1 | Alpha subunit of the oligosaccharyltransferase complex of the ER lumen | ✓ | ✗ |
| Pdi1 | Putative protein disulfide-isomerase | ✓ | ✓ |
| Pga63 | Component COPII vesicle coat; required for vesicle formation in ER to Golgi transport | ✓ | ✗ |
| Rot2 | Alpha-glucosidase II, catalytic subunit, required for N-linked protein glycosylation and normal cell wall synthesis | ✓ | ✓ |
| Sar1 | Functional homolog of *S. cerevisiae* Sar1; which is required for ER-to-Golgi protein transport; binds GTP; similar to small GTPase superfamily proteins | ✓ | ✗ |
| Sec13 | Putative protein transport factor | ✓ | ✗ |
| Sec23 | Putative GTPase-activating protein; regulated upon yeast-hypha switch | ✓ | ✗ |
| Sec24 | Protein with a possible role in ER to Golgi transport | ✓ | ✗ |
| Sec61 | ER protein-translocation complex subunit | ✓ | ✗ |
| Sec62 | Putative endoplasmic reticulum (ER) protein-translocation complex subunit | ✓ | ✗ |
| Ssh1 | Protein with a role in protein translocation across membranes | ✓ | ✗ |
| Stt3 | Putative oligosaccharyltransferase complex component | ✓ | ✗ |
| Wbp1 | Putative oligosaccharyltransferase subunit | ✓ | ✗ |
|  | Endocytosis: map04144 |  |  |
| Age3 | Putative ADP-ribosylation factor GTPase activating protein, functional ortholog of *S. cerevisiae* GCS1; mutation affects endocytosis | ✓ | ✗ |
| Cdc42 | Rho-type GTPase; required for budding and maintenance of hyphal growth | ✓ | ✓ |
| Chc1 | Clathrin heavy chain; subunit of the major coat protein; role in intracellular protein transport and endocytosis | ✓ | ✗ |
| Gea2 | Putative ARF GTP/GDP exchange factor | ✓ | ✗ |
| Glo3 | Putative ARF GTPase activator; role in COPI coating of Golgi vesicle, ER to Golgi vesicle-mediated transport, retrograde Golgi to ER vesicle-mediated transport | ✓ | ✗ |
| Hse1 | ESCRT-0 complex subunit; SH3-domain-containing protein | ✓ | ✗ |
| Pep8 | Protein similar to *S. cerevisiae* Pep8p, which is involved in retrograde transport | ✓ | ✗ |
| Rho1 | Small GTPase of Rho family; regulates beta-1,3-glucan synthesis activity and binds Gsc1p | ✓ | ✓ |
| Rim20 | Protein involved in the pH response pathway | ✓ | ✗ |
| Rsp5 | Putative NEDD4 family E3 ubiquitin ligase | ✓ | ✗ |
| Sec4 | Small GTPase of Rab family; role in post-Golgi secretion | ✓ | ✗ |
| Snx4 | Putative sorting nexin | ✓ | ✗ |
| Vps4 | AAA-ATPase involved in transport from MVB to the vacuole and ESCRT-III complex disassembly | ✓ | ✗ |
| Vps21 | Late endosomal Rab small monomeric GTPase involved in transport of endocytosed proteins to the vacuole; involved in filamentous growth and virulence | ✓ | ✗ |
| Vps27 | Putative ESCRT-0 complex protein with a role in multivesicular body (MVB) trafficking | ✓ | ✗ |
| Vps35 | Putative role in vacuolar sorting | ✓ | ✗ |
| Ypt31 | Protein required for resistance to toxic ergosterol analog | ✓ |  |
| Ypt52 | Rab-family GTPase involved in vacuolar trafficking, colocolizes with Vps1p and Ypt53p in late endosome | ✓ | ✗ |

We identified most of the proteins of the COPII vesicle component. Although the GEF protein Sec12 was identified in only one of the biological replicates of HEVs, Sar1, which is activated by Sec12 through phosphorylation, and Sec23/Sec24 and Sec13/Sec31 (Pga64) of the inner and outer coat complexes, respectively, were identified in all the biological replicates with significant q-values, while none of these proteins was identified in any of the YEV replicates. We also identified many proteins of the COPI complex (Sec26, Sec27, orf19.5689, Ret2, Sec21) plus the Arf1 protein and its activator GEF protein Glo3 (Table 1). This COPI complex is responsible for retrograde vesicle trafficking from the Golgi to the ER.

Looking into other intracellular protein trafficking pathways such as protein processing in the ER, endocytosis, and SNARE interactions in vesicular transport, it is remarkable how many proteins related to these processes were exclusively identified in HEVs protein cargo. Of the 19 *C. albicans* proteins represented in the KEGG vesicular transport pathway, 15 (78.9%) were identified in HEVs and not in YEVs (Figure S4). Similarly, conspicuous is the great number (32 out of 49) of the ER protein processing pathway identified in HEV cargo.

Only a small number of these proteins (Rot2, Kar2, Lhs1, Pdi1, orf19. 7578, Hsp70, Hsp90 and Msi3) were also identified in YEVs (Figure 5). When analyzing the set of proteins identified in HEVs by means of a protein-protein interaction network (STRING) and taking into account exclusively active sources of interaction with the highest confidence, we again observed HEV enrichment with proteins of the aforementioned pathways such as aa biosynthesis, endocytosis, purine metabolism and protein processing in the ER, and identified many proteosomal and ribosomal proteins (Figure 6).

**Figure 5.**
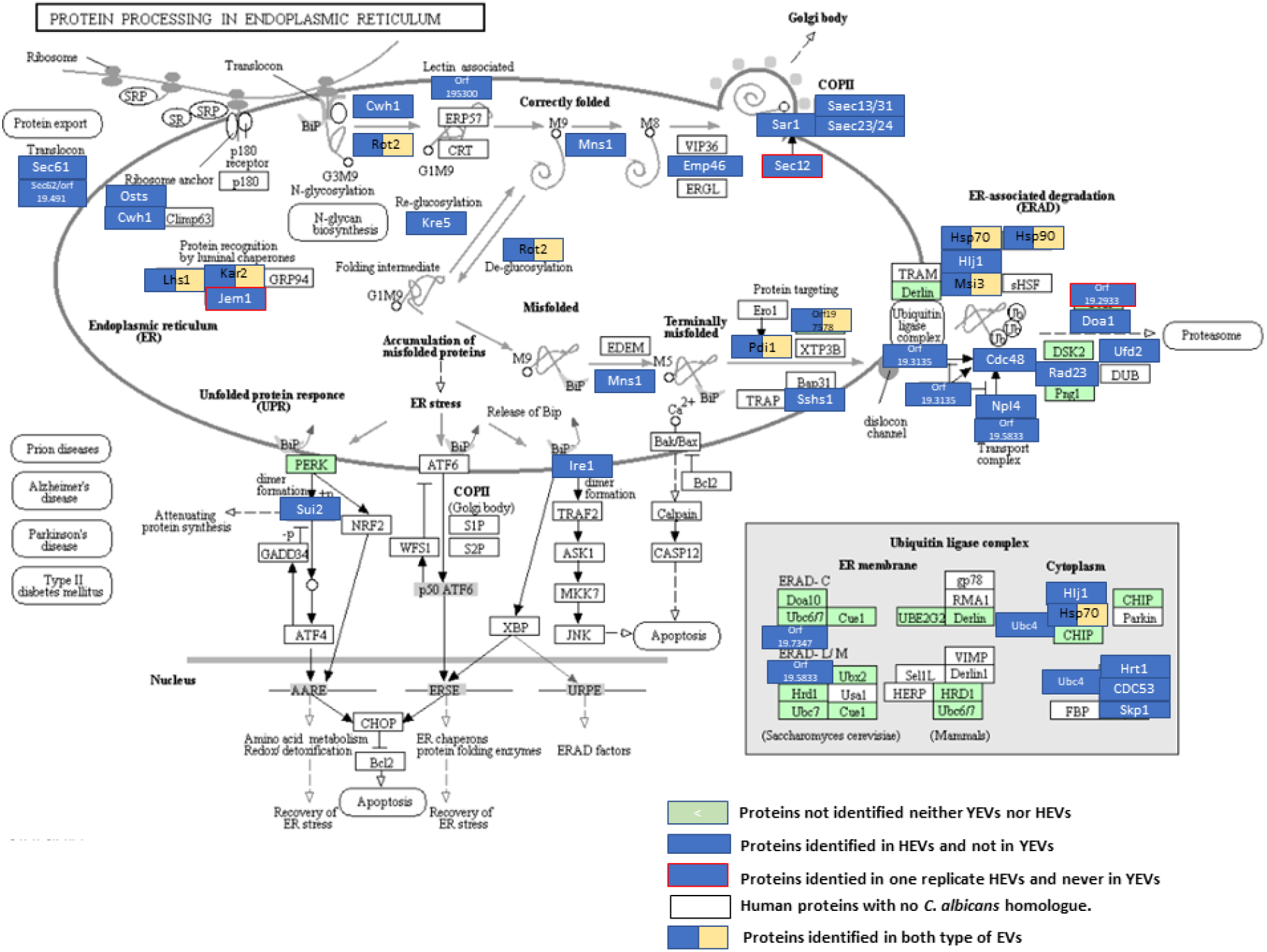
KEGG pathway related to processing in ER. Proteins exclusively identified in HEVs are filled in blue color. Those that have been identified in only one replicate are surrounded by red. Proteins identified in both types of EV are double colored (blue and beige).

**Figure 6.**
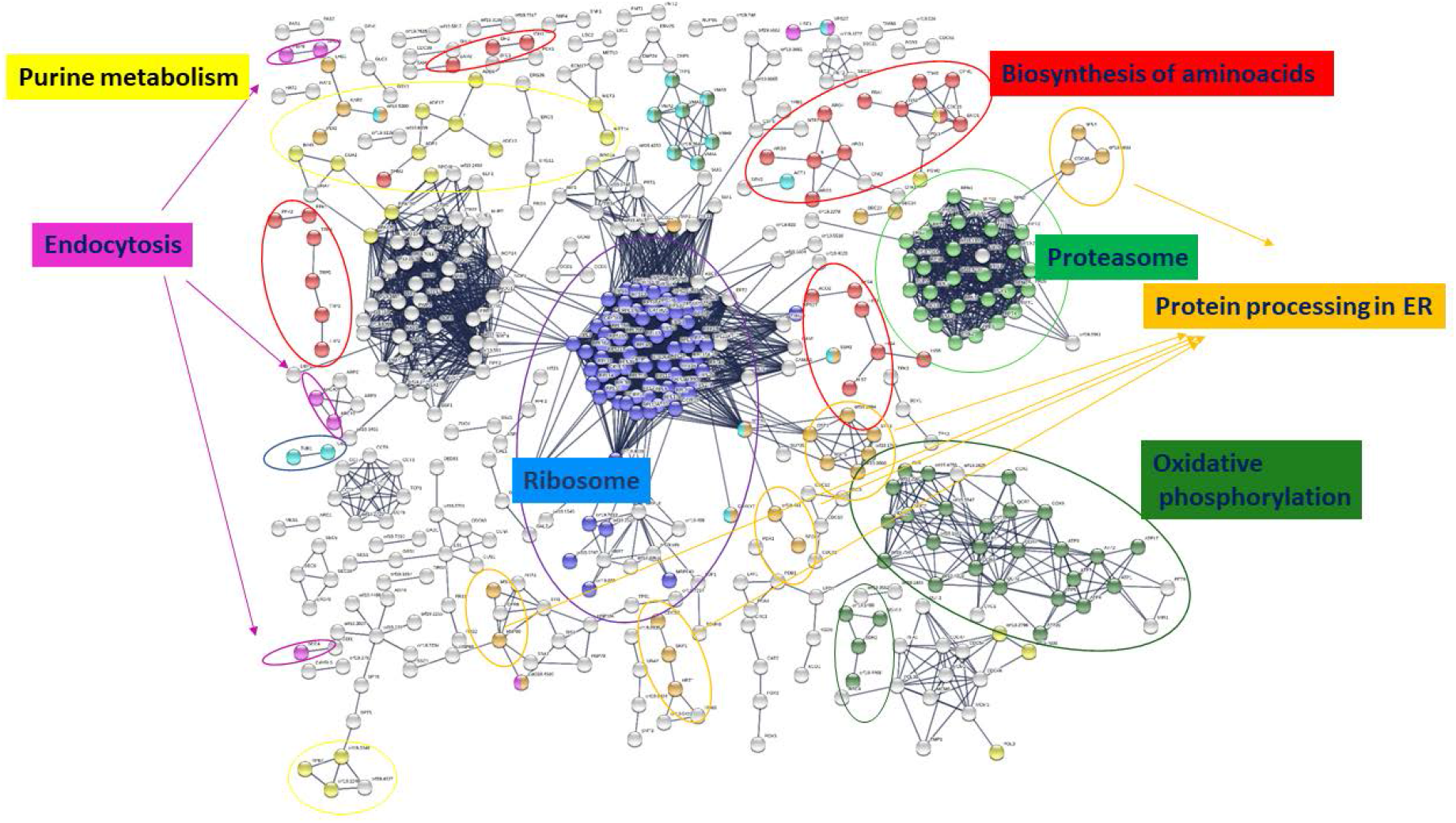
Protein-protein interaction network of proteins identified in HEVs. Only nodes corresponding to proteins with the highest confidence (0.900) in active interaction sources of co-ocurrence, co-expression, experiments, and neighborhood are shown.

#### 2. 4. Identification of virulence factors in *C. albicans* EVs

Lipases, phospholipases (PLBs) and secreted aspartic proteases (Saps) are classical virulence factors secreted by *C. albicans*. We identified several proteins with phospholipase activity in both types of EV. Nevertheless, while phospholipase B1 (PLB1), was exclusively identified in high amount in YEVs, PLB5, a putative GPI-linked phospholipase B and Plc2, a phosphatidylinositol (PtdIns)-specific phospholipase C (PI-PLC), were exclusively identified in HEVs.

Of the *C. albicans* Saps, we identified all but Sap 1 and Sap 3 in HEVs. Sap3 was identified exclusively in YEVs. Besides Sap3, four other Saps (Sap5, Sap7, Sap9, and Sap10) were identified in YEVs.

An interesting result regarding virulence factors is the identification of Ece1p protein in HEVs and not in YEVs. Ece1 proteolytic processing and maturation by Kex1-Kex2 proteins yield the *C. albicans* toxin candidalysin. Ece1p and both Kex proteins were identified in the protein cargo of HEVs, but the peptide corresponding to candidalysin was not. On the other hand, up to 16 possible predicted substrates for Kex2 were also identified in HEVs while only 7 were identified in YEVs.

Agglutinin-like sequence (Als) proteins are surface glycoproteins that mediate cell-cell aggregation and adherence to different surfaces. They are considered *C. albicans* virulence factors since they are adhesins. We identified Als3 in both types of EV, although more abundantly in HEVs (Figure 2). Als1p was also identified in HEVs, while Als2p was identified only in YEVs.

### 3. HEV enrichment in proteins, GO components and biological processes is specific and differs from that of hyphal whole-cell lysates (HWCLs)

Since we identified a much larger number of proteins in HEVs than in YEVs, we carried out a proteomic analysis of whole-cell lysates (WCLs) of cells of both morphologies in order to decipher whether this larger number of proteins in HEVs was due merely to a higher content of those proteins in the cytoplasm of hyphae. Considering the same parameters of being identified in at least 2 replicates with at least 2 peptides in a replicate, we identified up to 976 proteins and 587 proteins in yeast (YWCL) and hyphal (HWCL) WCL, respectively (Table S3).

The larger number and diversity of proteins in HEVs cargo were not due to their higher abundance in hyphal cytoplasm, since the majority of proteins identified in HEVs cargo were not identified in HWCL. In the case of yeast cultures, we identified many more proteins in YWCL than in YEVs.

Comparison of GO enrichment analyses of proteins from HEVs and HWCL confirmed that the most significant GO cellular component and biological processes were differed between the protein extracts. HEVs, but not HCWL, were enriched with proteins from the plasma membrane, ER membrane, and endosome membrane in the GO cellular component analysis, suggesting the involvement of these endomembrane systems in the biogenesis of HEVs (Figure 7). Similarly, HEVs but not HWCL were enriched with enzymes from the *de novo* purine and pyrimidine biosynthetic pathways as well as processes related to the formation of cytoplasmic translation initiation complexes and the cellular response to oxidative stress (Figure S5). Also interesting was the HEV enrichment with the ergosterol biosynthetic process that was not observed in HWCL. Thirteen proteins from ergosterol biosynthesis, including Erg11 (the main target of azoles),^36^ were identified in the protein cargo of HEVs while only three (Erg10, Erg 13 and Erg 20), were identified in YEVs, with lower NSAF values (TableS4). Furthermore, the HEV proteins also included nine described as required for resistance to toxic ergosterol analog ^37^(Table S4). Moreover, 127 proteins identified in HEVs have been described as either induced by azole treatment or linked to azole resistance (Table S4).

**Figure 7.**
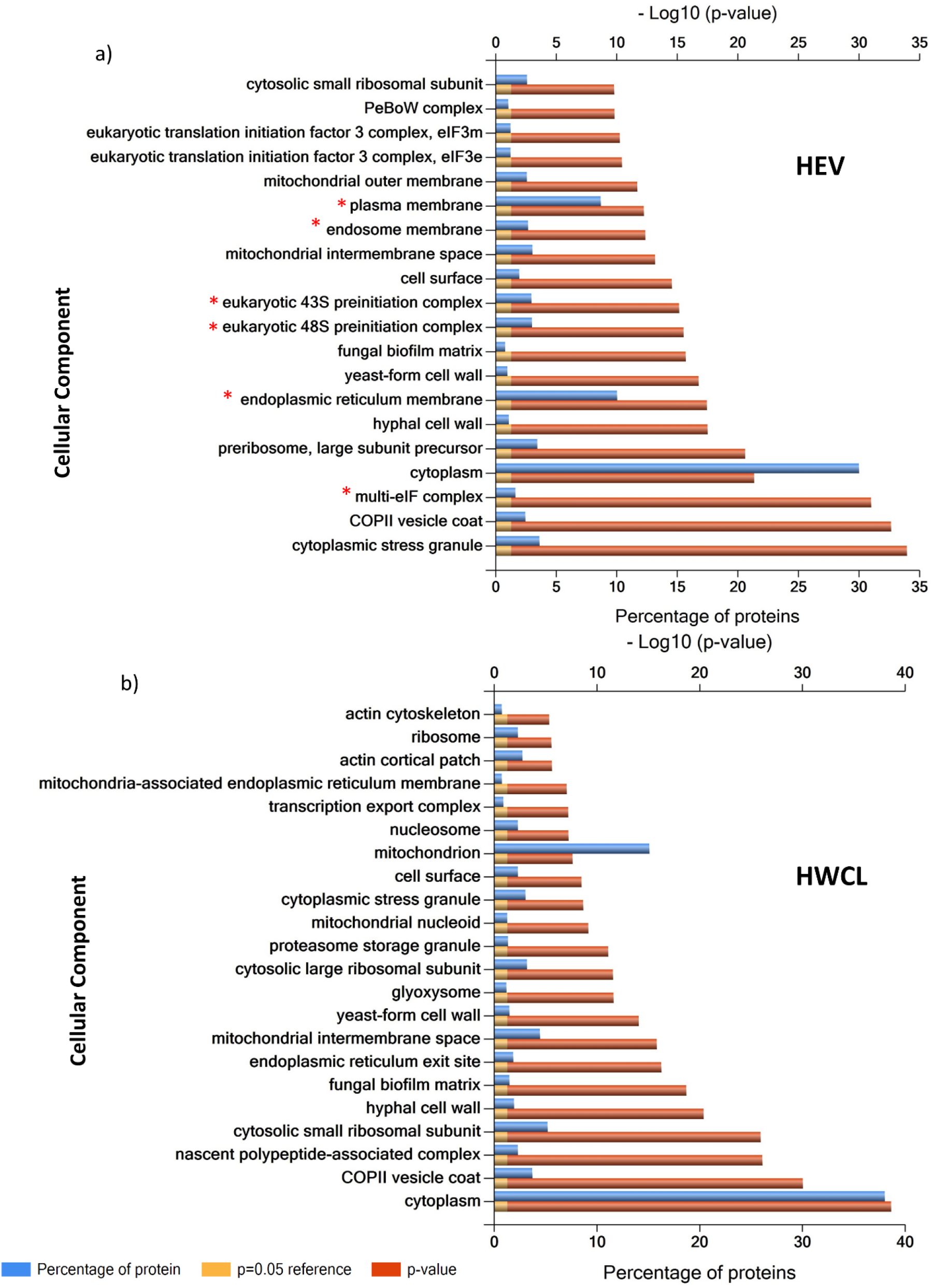
FunRich categorization of component enrichment of proteins identified in a) HEVs and b) HWCL (Hyphal Whole Cell Lysate). *Cellular components exclusively enriched in HEVs.

### 4. YEVs and HEVs contain immunoreactive proteins but differs in the induction of TNFα release from THP1 macrophages

Many cell surface proteins from *C. albicans* are reported to be immunogenic. We tested proteins present in YEVs and HEVs for reactivity with human sera from invasive candidiasis patients since both types of EVs contained large amounts of cell wall and surface proteins (Figures 2 and 3a). Usually, these cell wall proteins have high molecular weights because they are highly glycosylated. Both types of EV contained these kinds of proteins although their electrophoretic pattern were clearly different (Figure 8a). Figure 8b shows that proteins from both types of EV proteins were recognized by the sera but with different recognition patterns depending on the cell morphology of origin. Human immunoglobulins from patients with invasive candidiasis recognized high-molecular weight proteins with a strong signal in YEVs proteins. On the other hand, there were other proteins of lower molecular weights detected exclusively in HEVs.

**Figure 8.**
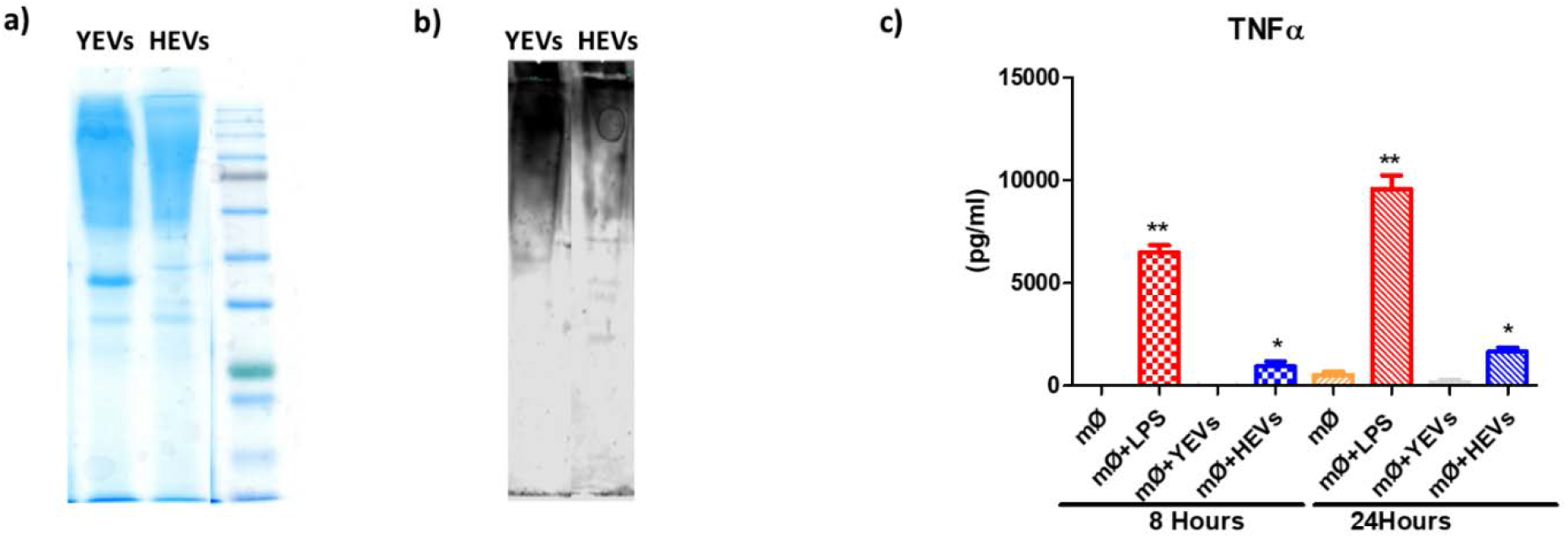
a) SDS-PAGE Coomasie-blue stained gel of YEVs and HEVs protein extracts. b) Western blot showing the immuno-reactive pattern of YEVs and HEVs protein extracts probed with sera from patients suffering invasive candidiasis. c) TNFα release from TPH1 activated macrophages incubated during 8 or 24 hs with 5µg of YEVs or HEVs. Negative and positive controls (by addition of PBS and LPS respectively) were also analysed. * t-test p value <0.02; ** t-test p value <0.001

Regarding the ability to stimulate the secretion of cytokines from THP1 derived macrophages, only HEVs were able to increase the secretion of TNFα at the two time points assayed, increasing the amount of this released cytokine at 24 hs (Figure 8c). No significant differences were observed in the case of the secretion of other cytokines, such as IL-10 and IL-12 (data not shown). We also tested both types of EVs for their ability to damage macrophages, by incubating 5µg of HEVs and YEVs with macrophages for 4 or 8 hs and measuring the LDH activity in the culture supernatant. Since LDH is an internal enzyme, increases in the abundance of this enzyme in culture supernatant would indicate cell damage. Although an increasing tendency in cytotoxicity effect was observed in the case of HEVs, it was not statistically significantly different from the YEV or negative control results (data not shown).

## DISCUSSION

The study of EVs has increased dramatically during recent years due to the high number of physiological processes in which these membranous structures are involved.^23, 28, 38^ Since the *C. albicans* dimorphic transition is one of the main virulence traits of this fungus, the study of EVs secreted by cells of each type of morphology could be highly relevant to the battle against invasive candidiasis and spur the possible discovery of putative new antifungal targets. We have obtained YEVs and HEVs from *C. albicans* by ultracentrifugation and found interesting differences between them, in both physical properties and protein cargo.

Different sizes of EVs has been described in different works, and both the specific strain and the culture conditions seem to contribute to the wide heterogeneity in size, protein cargo, and biogenesis of EVs.^10^ In our culture conditions, HEVs were in general smaller than YEVs, with a larger population of vesicles in the size range of 100-200 nm, in contrast to 400-500nm for YEVs. In accord with our results, a recent analysis of EVs secreted by biofilm and yeasts from reference *C. albicans* strain DAY286 revealed that EVs secreted by biofilms are also of smaller size.^39^

The differences observed regarding protein cargo are also very relevant. We identified six times more proteins in HEVs than in YEVS, analyzing the same amount of protein from each. Proteins common to both YEVs and HEVs represented 92% of the proteins in YEVs but less than 16% in HEVs. This fact and the DLS data strongly suggest that the collected HEVs were probably a mixture of EVs types, with a proportion similar in size and protein cargo to YEVs in addition to other, different EVs, which were smaller and contained more than 1000 proteins not identified in YEVs. These differences are probably related to their different mechanisms of biogenesis.

The disparity in the number of proteins identified depending on cell morphology has been also observed in other *C. albicans* studies of extracellular proteins, such as that by Luo et al. who studied the yeast-to-hypha transition and identified four times more proteins in the hyphal than in the yeast secretome.^32^ Similarly, in a study of the cell surface proteins (the surfome) Gil Bona and coworkers used tryptic digestion of live yeast and hyphal *C. albicans* cells to describe around 400 and almost 900 proteins in yeast and hyphal cells, respectively.^7^ All these data are evidence that not only the cell surface but also the extracellular environment (secreted proteins and EVs) have higher protein diversity in hyphae that in yeast. It is plausible that this higher protein diversity contributes to hyphal adhesion and tissue invasion.

### Differences in virulence factors and immunomodulatory effect between YEVs and HEVs

The exclusive presence in HEVs of many proteins already described as virulence factors, such as Ece1p, is remarkable. Ece1p is the candidalysin preproprotein, a fungal peptide toxin critical for mucosal infection.^40, 41^ After sequential proteolytic processing by Kex2, a Golgi-located subtilisin-like protease, and the carboxypeptidase Kex1 candidalysin is secreted and can be detected in culture supernatants and during growth on epithelial cells.^40,42, 43^ We identified 13 different peptides from Ece1p covering 63% of the protein sequence, and both Kex2 and Kex1, though not the peptide corresponding to candidalysin. This fact does not necessarily support the lack of the toxin in HEVs because, due its structure, it could be inserted into the membrane vesicles and might be more difficult to extract or and detect. In fact, in the peptide atlas of *C. albicans*, this peptide is described as having been identified only in samples obtained by shaving the cell surface.^44, 45^ More experiments are needed to decipher the real presence of candidalysin in HEVs, which would represent an alternative mode of Ece1 processing and secretion. Although candidalysin has been described as promoting cell damage, the minimum concentration to elicit the damage was 5µM of pure toxin.^46^ Thus, it is not surprising that even though our HEVs might contain this toxin we did not observe damage in our macrophage HEV assays, since we used a much smaller amount (5 µg of HEVs).

Other virulence factors secreted by *C. albicans* such as Saps, Als, and PLBs were also identified in both types of vesicles, although with marked differences in the type and abundance depending on the morphology of origin. Sap proteins, comprising Sap1 through Sap10, participate in host tissue and protein degradation, contributing to invasion and nutrient uptake.^47^ We identified all but Sap1 and Sap3 in HEVs. Sap3 was identified exclusively in YEVs. Besides Sap3, four other Saps (Sap5, Sap7, Sap9, and Sap10) were identified in YEVs.

The greater abundance of Sap 5 in HEVs, together with the identification of Sap 4 and Sap 6 exclusively in HEVs, are in line with the fact they were only identified only in the secretome of hyphae-enriched cultures grown in the presence of N-acetyl glucosamine at pH7.4 and not in pure yeast cultures.^31^ The presence of these Saps (4, 5, and 6) is reportedly essential for *C. albicans* systemic invasion.^48^ Sap7, Sap9, and Sap10 had higher NSAF values in YEVs than in HEVs (Figure 2). According to Albrecht et al, Saps 1-Sap8 are secreted, while Sap9 and Sap10 are retained at the cell surface via a modified GPI anchor.^49^ Since YEVs are enriched in cell surface proteins, the higher abundance of these GPI-linked Saps within YEV cargo is not surprising. This fact is also in agreement with the hypothesis proposed by Gil-Bona et al., and commented on below, of YEVs’ main origin in outward budding from the plasma membrane.^19^ In agreement with this hypothesis, although in general the abundance of all common phospholipases was higher in YEVs, the highest difference in abundance was observed for Plb4.5 (identified in YEVs with an NSAF value 10-folg greater than in HEVs), which contains a GPI anchor and it is likely to be retained at the cell surface. This phospholipase was also identified exclusively in the secretome from yeast-form.^31^ On the other hand, Plb5, a putative GPI-linked phospholipase B that has been described as fungal-specific (with no mammalian homolog) and whose null mutation attenuates virulence, was exclusively identified in HEVs. Phospholipase C (PI-PLC), another phospholipase identified exclusively in HEVs, is reportedly induced under filamentation conditions.^50^

The Als proteins Als1 and Als2 were, respectively, identified in HEVs and YEVs exclusively. Als3, which has been directly linked to adhesion to and invasion of human cells was more abundant in HEVs than in YEVs (Figure 2).^51^ Als3, together with Ssa1 (also identified in both types of EV), has been described as being linked to E-cadherin and promoting induced endocytosis.^52^ Furthermore, Mp65, a β-glucanase directly implicated in *C. albicans* adhesion as a mediator of attachment to various substrates was very abundant in both types of EV. Considering that the initial adhesion of *C. albicans* cells to epithelial cells likely occurs between yeasts and epithelial cells, the high Mp65 content of YEVs is not surprising.^53^

The immunomodulatory effect of EVs secreted by different organisms, including *C. albicans* has been recently described.^20, 28^ Cell surface proteins have been proven to be immunogenic in many works.^54^ The presence in both types of EVs of many immunogenic proteins was demonstrated by the use of western blots probed with sera from a patient suffering invasive candidiasis (Figure 8b). Nevertheless, the signal corresponding to highly glycosylated proteins, which are expected to be mainly cell surface proteins, was stronger in YEVs (Figure 8b). This is in accord with the fact that cell surface proteins, although being identified in both types of EVs, were more abundant in YEVs according to the calculated NSAF values (Figure 2). However, in the conditions tested, only HEVs were able to enhance the induction of TNFα cytokine when incubated with THP1 macrophages. Even though the release of TNFα by bone marrow-derived murine macrophages stimulated by *C. albicans* YEVs has been described, it is worth mentioning that the *C. albicans* strain used in that study was a clinical isolate and YEVs were obtained after 48 h incubation in Sabouraud medium.^20^ The *C. albicans* strain, time of EV collection, culture conditions, and the immune cell origin were all completely different from ours. In our conditions, EVs isolated from cells of the two different morphologies had significantly different interactions with the THP1 macrophages. All these data point to the relevance of HEVs in *C. albicans* infection, in dissemination of virulence factors and other components via the vesicles or in interactions with immune cells.

### Proteins involved in protein metabolism in *C. albicans* EVs

It is important to highlight the large number of proteins involved in protein metabolism that were identified in HEVs but not in YEVs, such as the numerous proteasome complex proteins identified exclusively in HEVs. The proteasome is a multi-subunit protein structure comprising two 19S regulatory particles surrounding a catalytic core 20S particle. This complex degrades misfolded and damaged proteins in the cell and ultimately is critical in many cellular processes such as cell-cycle regulation, cell differentiation, signal transduction pathways, inflammatory responses, and apoptosis.^55, 56^ A recent work describes the important role the *C. albicans* proteasome plays in fungal morphogenesis. Inhibition of many proteins from both the catalytic and regulatory particles of the proteasome induced hypha formation.^57^ This work also describes the essentiality of all but five genes (*SEM1, RPN10, RPN13, RPN9*, and *PRE9*) encoding proteasomal proteins, identifying Sem1 as a proteosome protein. We identified in HEVs all but five proteins (Rpn10, Rpn13, Rpn15, Pre6, and Pre10) of the proteasome complex besides Sem1. Furthermore, our group has analyzed the proteomic response of *C. albicans* to 10 mM hydrogen peroxide (H_2_O_2_) and observed an increase in the abundance of different proteasomal proteins from the catalytic subunit. This increase in protein abundance was subsequently corroborated functionally by measuring the catalytic activity of proteasomes in the presence of 10 mM H_2_O_2_ (unpublished results). Since *C. albicans* faces oxidative conditions in its battle against the human immune system, it could be advantageous for it to shuttle large amounts of this complex in HEVs. The presence of proteins from this complex in EVs has also been reported in other microorganisms, such as *Acanthamoeba castellani*. Thirteen proteasome subunits were identified in exosome-like vesicles secreted by this organism^58^ which can cause severe keratitis in healthy individuals (particularly contact lens users) and amoebic encephalitis, disseminated disease, or skin lesions in individuals with compromised immune systems.

Beside proteasome complex proteins, the enrichment of HEVs in proteins related to protein synthesis, including proteins from ribosomes, the translation initiation complex, and aa biosynthetic pathways, suggests that HEVs are somehow involved in *C. albicans* protein metabolism. In fact, since HEVs contain ribosomal proteins, most of the proteins from the preinitiation multi-eIF complex, and most of the enzymes necessary for most aa biosynthesis, it would be possible that HEVs synthesize and are a reservoir of aa.

Also remarkable is the identification in HEVs of all the enzymes involved in the conversion of PRPP to IMP and then to AMP, while only three of them were identified in YEVs. Apart from those proteins, Imh3 and Gua1, which are involved in the synthesis of GMP from IMP, were also exclusively identified in HEVs. Curiously, mutations in the *ADE8* and *GUA1* genes are related to lack of virulence, since mice infected with these deletion mutants showed 100% survival and undetectable fungal kidney loading.^59^ Purines are essential molecules not only in DNA and RNA backbones, but also in many metabolic pathways, including energy utilization, regulation of enzyme activity, and cell signaling. Hence, it is not strange that all the enzymes needed to synthesize this valuable resource are packaged in EVs to be secreted and shared by all *Candida* cells in the community.

### *C. albicans* HEVs and YEVs could differ in their biogenesis

Apart from cell surface proteins also contained in YEVs, HEVs contain numerous cytoplasmic proteins. Even though HEVs contained 88% of the proteins present in YEVs, the 100 most abundant HEV proteins are cytoplasmic, in contrast to YEVs in which the 100 most abundant proteins are cell surface related. This piece of evidence, together with the existence of a small proportion of larger vesicles secreted by hyphae (more similar in size to YEVs) suggests that hyphae might produce two different types of EV: bigger HEVs whose protein cargo are probably cell surface and membrane-related proteins commonly identified in YEVS, and a larger proportion of smaller HEVs enriched in cytoplasmic proteins, with the two types produced by different mechanisms. In human cells, depending on their mode of formation, EVs are generally separated into two categories: exosomes of endolysosomal origin that are released upon fusion of multi-vesicular bodies (MVB) with the plasma membrane and ectosomes, which are microvesicles that form through cell surface budding. EV size has been one of the most widely used criteria for vesicle classification, with small (<150 nm) vesicles classified as exosomes, while ectosomes are larger (100 to >1000 nm).^60,61, 62^ ESCRT machinery has been linked to exosome biogenesis. ESCRT components have been identified in exosomes in many proteomic studies (see EVpedia: www.evpedia.info; Vesiclepedia: www.microvesicles.org). In *C. albicans*, ESCRT pathway-related mutants were deficient in vesicle secretion from biofilms (a morphology intrinsically related to hyphal forms) associating this pathway with vesicle secretion.^26^

Our data agree with the proposed biogenesis mechanisms. Based on the bigger size and the enrichment in cell wall and cytoplasmic membrane proteins of the majority of YEVs, these vesicles are more likely to be microvesicles budded, pinched off, and released to the extracellular space from the plasma membrane, as proposed.^19^ This is probably also the origin of the larger HEVs. In contrast, the contents of the larger proportion of smaller HEVs, including many proteins from different endomembrane compartments such as the ER, COPII vesicles, endosomes, MVB, and vacuole, including ESCRT components, suggests their origin in these protein-trafficking regions of the cell and exosomal nature. Consistent with this notion, we have identified exclusively in HEVs proteins such as Sec4p, whose absence alters the composition of vesicles.^63^ Other proteins related to MVB formation have also been exclusively identified in HEVs, as have proteins belonging to the ESCRT pathway (Vps4 from ESCRTIII, and Hse1 and Vps27 from ESCRT-0) (Table 1). The presence in HEVs of most of the proteins from the COPI retrograde Golgi-to-ER vesicle-mediated transport complex and COPII anterograde ER-to-Golgi vesicle-mediated transport complex (Table 1) supports the intracellular origin of HEVs.

A recent study by Dawson et al. defined specific protein markers for *C. albicans* EVs.^39^ The authors proposed a list of 42 proteins found to be exclusive to or significantly concentrated in EVs across all the strains assayed, which included yeast forms from two clinical isolates and yeast cells and biofilm from the laboratory strain DAY286. They considered Sur7p and Ewp1 (orf19.6741) to be the best marker candidates for *C. albicans* EVs, since these proteins were present in all EVs from all strains and morphologies assayed and shared topological features with tetraspanins, which are the accepted markers for human exosomes. We identified these two proteins among the proteins identified in both YEVs and HEVs, but also identified in both types of vesicle some proteins described by Dawson et al. as “negative EV markers” since they could not identify these proteins in any of their EV protein extracts. These results confirm the high variability in EV protein loading depending on strain and culture conditions and described in numerous works.^10^ Thus, considering the high heterogeneity in kind and size of EVs reported in *C. albicans*, further proteomic studies are required before positive and negative *C. albicans* EV markers can be properly defined and confirmed.

## CONCLUDING REMARKS

This work represents the first analysis of EVs from *C. albicans* hyphae and their comparison with EVs from the yeast form. The data obtained from EVs collected by ultracentrifugation reveals a higher proportion of smaller EVs in hyphal than in yeast cultures. These HEVs carry a large number of proteins not present in YEVs, including not only virulence factors but also many proteins involved in central cellular functions such as protein, purine, and pyrimidine metabolism and vesicular trafficking, the latter comprising several proteins related to exosome biogenesis.

The differences between YEVs and HEVs also prompted a different immune host response, as tested with in vitro macrophage cell cultures and human sera from patients with candidiasis. All these results show the relevance of EVs, and mainly HEVs, to *Candida* infection. More studies on HEVs to elucidate their biogenesis and their role in cell-cell communication and infection would be very interesting in relation to fighting invasive candidiasis. Considering the increasing number of biomedical applications proposed for exosomes, this knowledge could also help the development of new diagnostic tests and/or therapeutic strategies

## AUTHOR INFORMATION

### Author Contributions

R.M., E.R., G.C., M.L.H. and S.R. performed the experiments. R.M. contributed to the conception and planning of the study and wrote the manuscript. C.G. and L.M. Conceived and designed the experiments, supervised the experimental work and critically reviewed the manuscript.

## ACKNOWLEDGMENT

We gratefully acknowledge Alberto Bottos for his collaboration in the whole cell extracts preparation for the proteomic analysis. This work was supported by BIO2015-651472-R from the Ministry of Economy and Competitiveness, RTI2018-094004-B-100 from Spanish Ministry of Science and Innovation, InGEMICS-CM B2017/BMD3691 from the Comunidad de Madrid, Spanish Network for the Research in Infectious Diseases (REIPI RD16/0016/0011) and PRB3 (PT17/0019/0012) from the ISCIII. InGEMICSCM, REIPI and PRB3 are cofinanced by European Development Regional Fund ERDF “A way to achieve Europe”. The proteomic analyses were performed at the Proteomics facility of Complutense University of Madrid (UCM) a member of ProteoRed-ISCIII network and at Thermo Fisher Scientific GmbH, Dreieich, Germany. DLS was carried out at the Spectroscopy and correlation facility of the UCM. TEM was carried out at the Electronic microscopy facility of the UCM. RM was supported by InGEMICS-CM and RTI2018-094004-B-100. These results are lined up with the Human Infectious Diseases HPP initiative from the Human Proteome Project (HID-HPP).

## ASSOCIATED CONTENT

**Table S1**. Excel document with proteins identified in at least two biological replicates with at least two peptides in one replicate in HEVs and YEVs.

**Table S3**. Excel document with proteins identified in at least two biological replicates with at least two peptides in one replicate in HWCL and YWCL.

**Figure S1.**
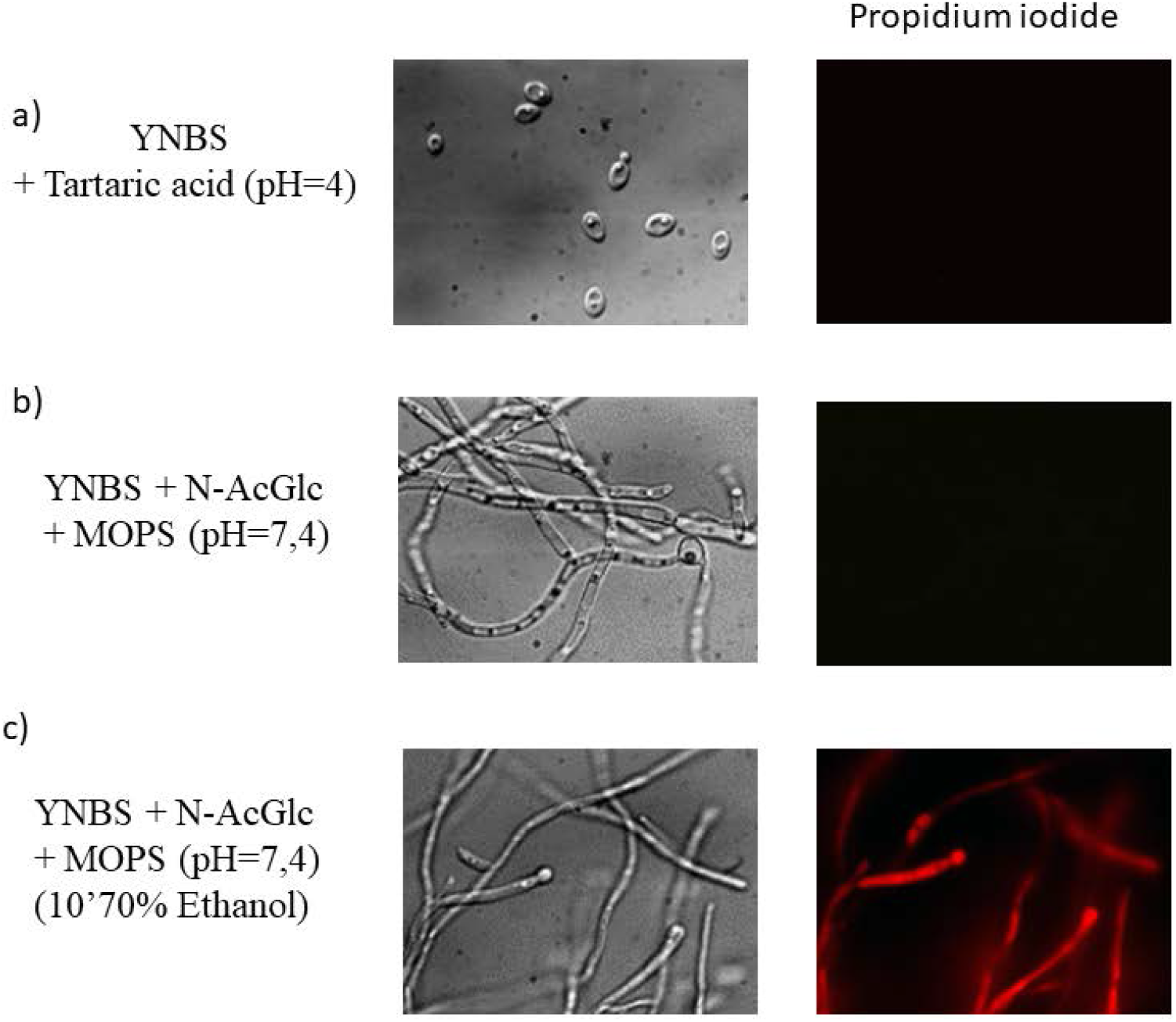
Phase-contrast micrographs and PI stained micrographs of *C. albicans* yeast and Hyphal cells.. a) yeast (YNBS + Tartaric acid (pH=4)) and b) hyphal (YNBS + N-AcGlc + MOPS (pH=7,4)) cultures were treated with propidium iodide. Phase-contrast micrographs and PI stained micrographs (λ=488nm) were acquired using a 40x oil objective. Positive PI control c) was obtained by ethanol 70% treatment for 10 minutes.

**Figure S2.**
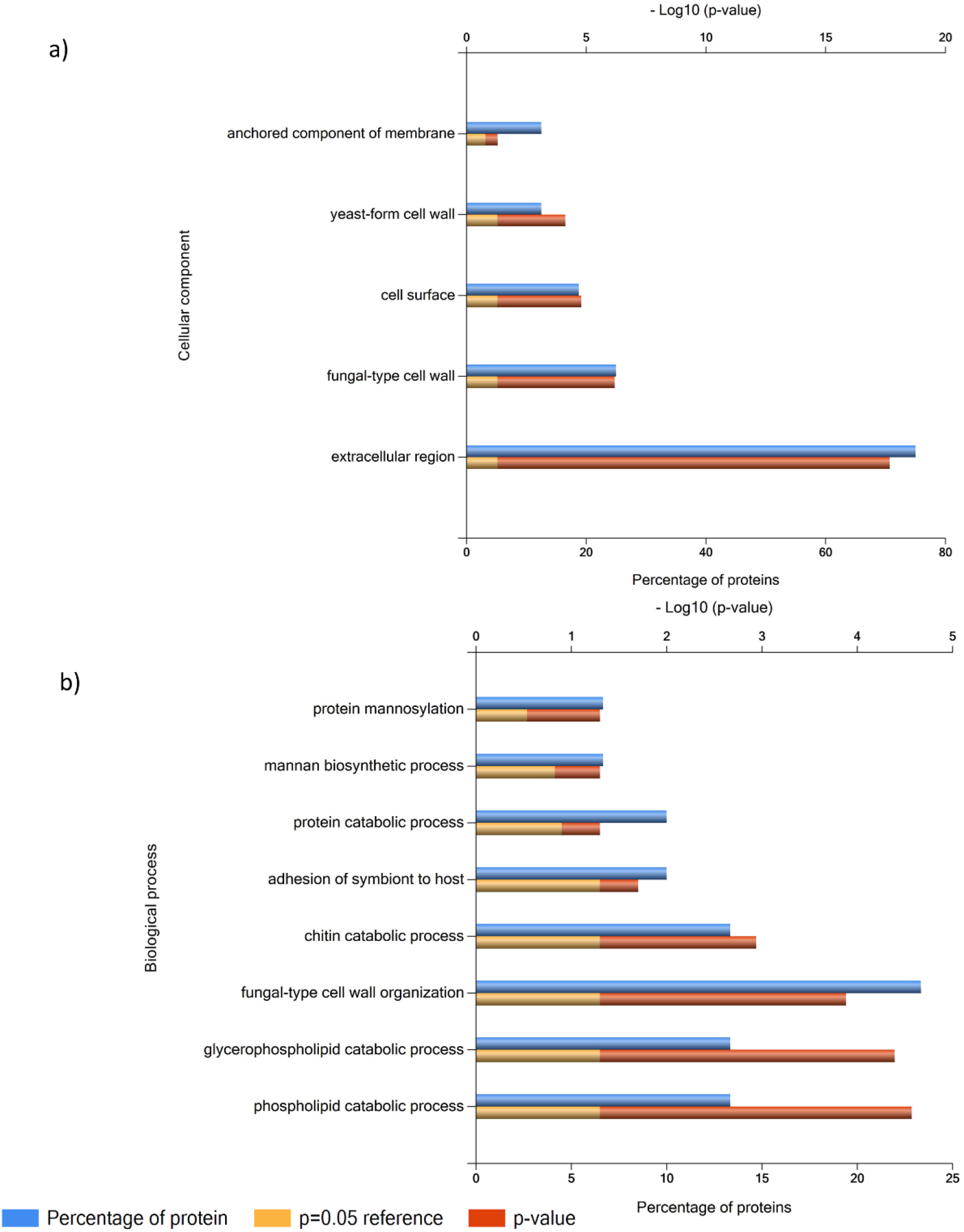
FunRich categorization of component a) and biological process b) enrichment of proteins exclusively identified or enriched in YEVs (Table S2)

**Figure S3.**
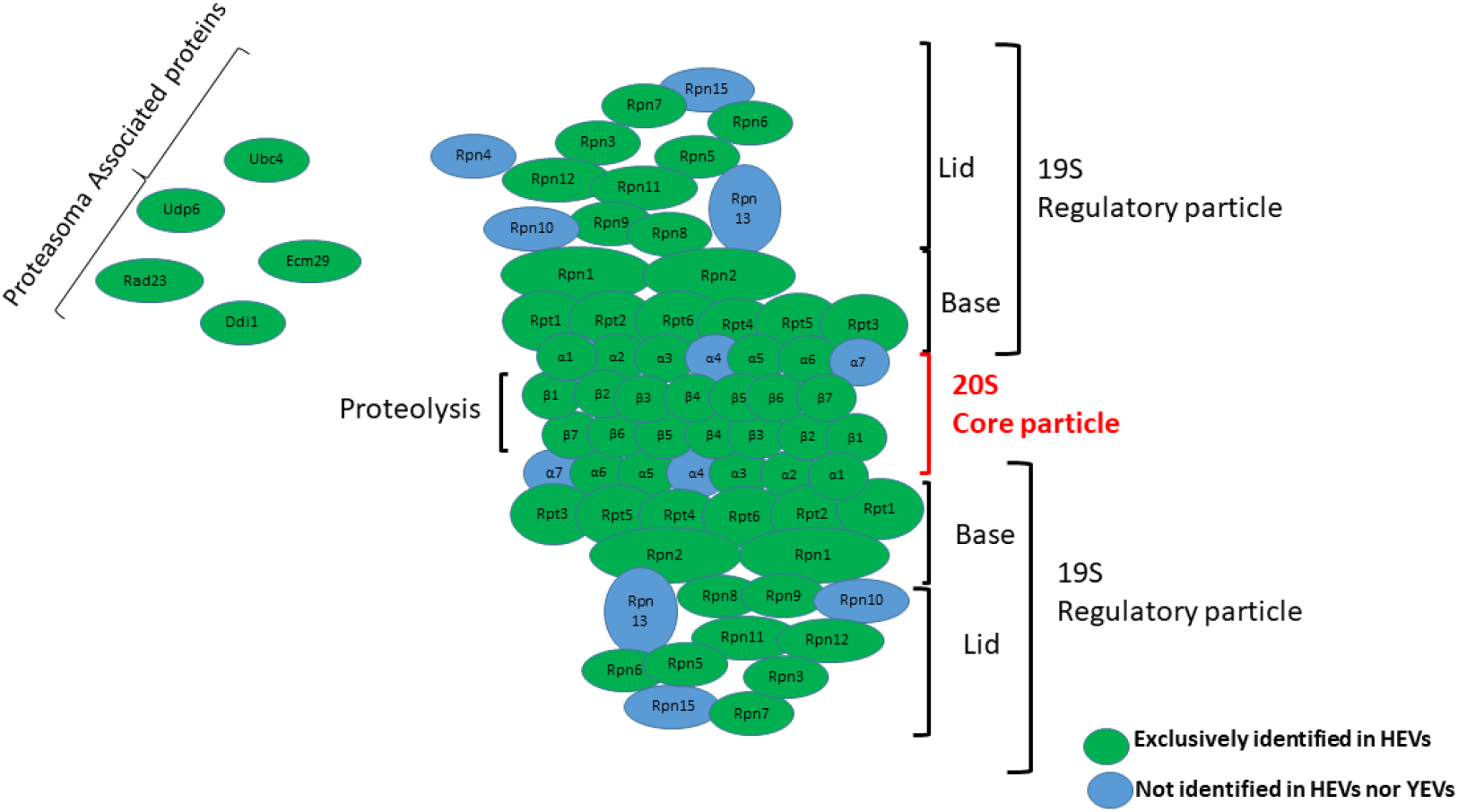
Schematic representation of the proteasome complex showing all the proteins from the 20S core particle and 19S regulatory particle.

**Figure S4.**
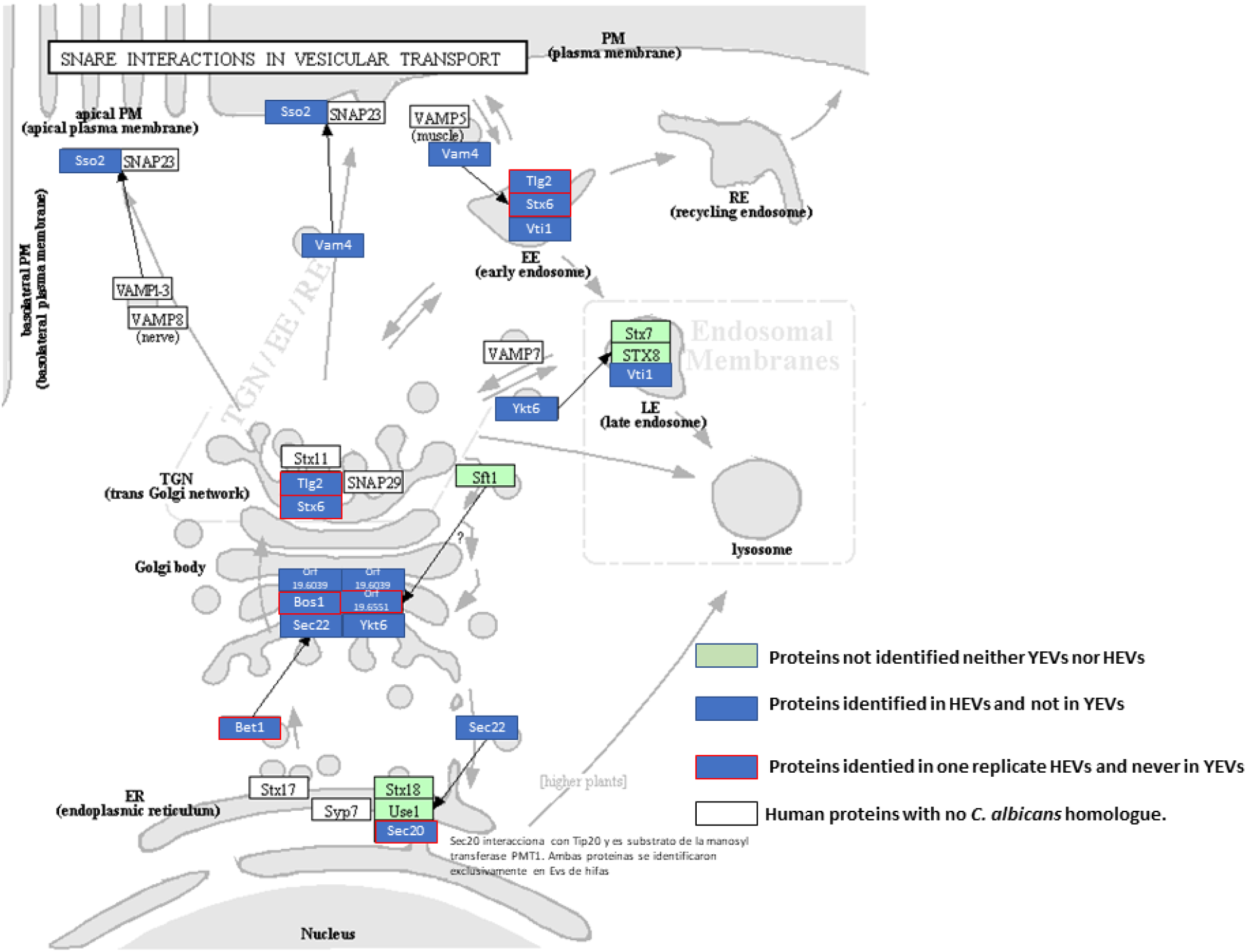
*C. albicans* proteins exclusively identified in HEVs related to SNARE process. *C. albicans* homologues to human proteins involved in SNARE process which have been exclusively identified in HEVs are filled in blue. Those only identified in one of the replicates are surrounded in red. Human proteins with no *C. albicans* homologue are in white.

**Figure S5.**
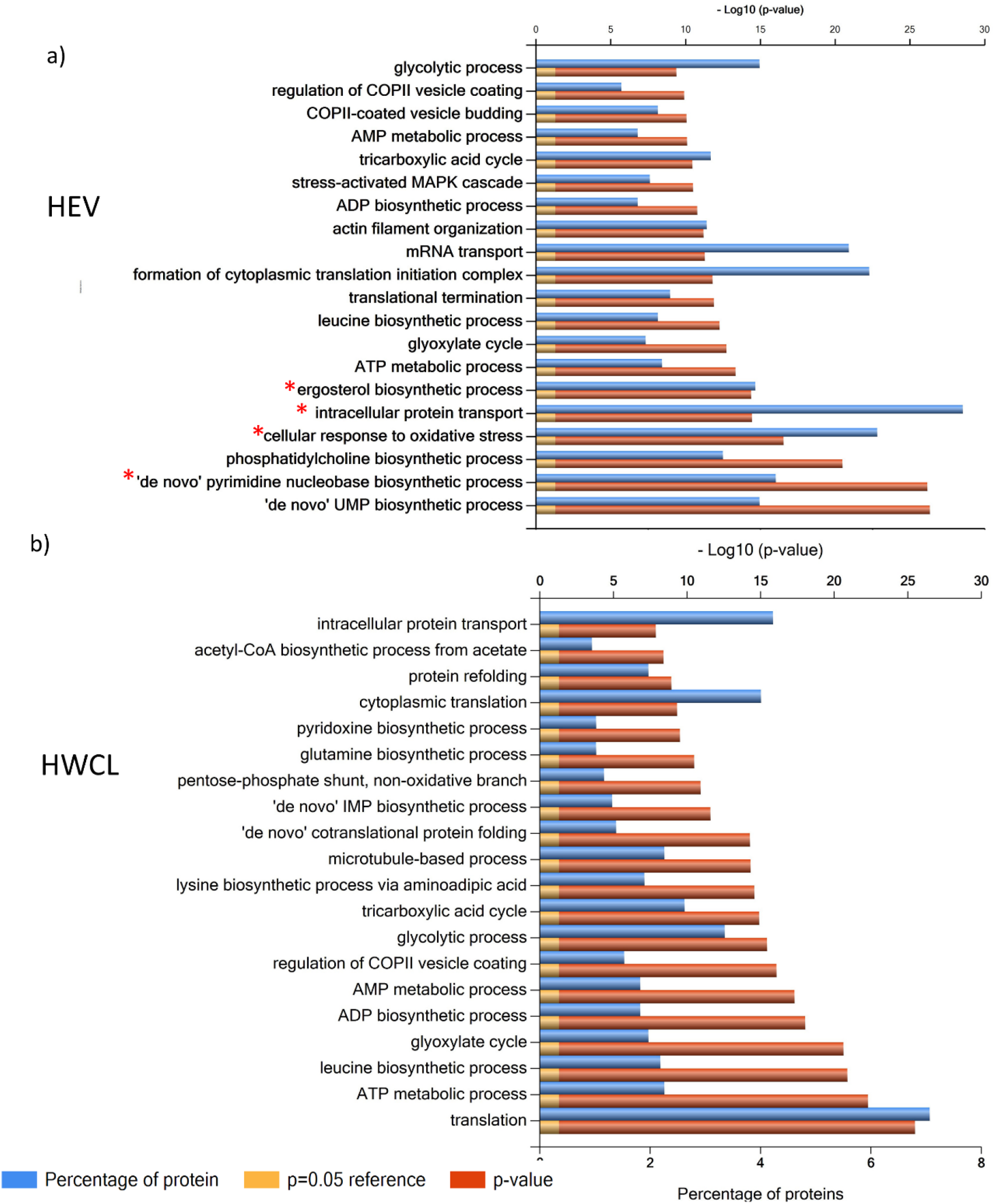
FunRich categorization of biological process enrichment of proteins identified in a) HEV and b) HWCL.

**Table S2.** List of proteins exclusively identified or enriched in YEVs (also identified in HEVs but with at least 10-fold lower NSAF values) ordered in decreasing NSAF value.

| Protein | NSAF YEVs | NSAF HEVs |
| --- | --- | --- |
| Rbe1 | 0,02157365 | 0,00019836 |
| Dag7 | 0,01198828 | 0,00019753 |
| Sap3 | 0,00762739 |  |
| Orf19.5322 | 0,00611279 | 0,00020397 |
| Orf19.2870 | 0,00437018 |  |
| Cht3 | 0,0043248 |  |
| Plb1 | 0,00382391 |  |
| Orf19.1766 | 0,00372026 | 0,00025655 |
| Ldg3 | 0,00357281 |  |
| Cht1 | 0,00342355 | 0,00023609 |
| Sap10 | 0,00339577 | 0,00016392 |
| Orf19.6119 | 0,00318245 | 0,00021436 |
| Orf19.6487 | 0,00300739 |  |
| Pir1 | 0,00252264 |  |
| Rbt7 | 0,00250207 |  |
| Orf19.6741 | 0,00242355 | 0,00012856 |
| Yps7 | 0,00228221 | 0,00019038 |
| Ywp1 | 0,00211066 |  |
| Fmp45 | 0,00178858 |  |
| Fet99 | 0,00172186 |  |
| Orf19.1765 | 0,00147473 | 0,00010983 |
| Hom2 | 0,00135239 |  |
| Mnn23 | 0,00124276 | 0,00021563 |
| Tfs1 | 0,00123797 |  |
| Orf19.540 | 0,00122646 |  |
| Fma1 | 0,00112015 | 0,00016854 |
| Gcy1 | 0,00109917 |  |
| Cht2 | 0,0010555 | 0,00013292 |
| Orf19.2484 | 0,00094184 | 0,00024356 |
| Orf19.2516 | 0,00091649 | 0,0001896 |
| Orf19.4211 | 0,00086946 |  |
| Glo1 | 0,00077702 | 0,00026304 |
| Msb2 | 0,00073218 | 0,00015966 |
| Hbr2 | 0,00060133 |  |
| Plb3 | 0,00060047 | 0,00014907 |
| Pho113 | 0,00059829 |  |
| Mnn22 | 0,00057277 |  |
| Gcv3 | 0,00055804 |  |

**Table S4.**
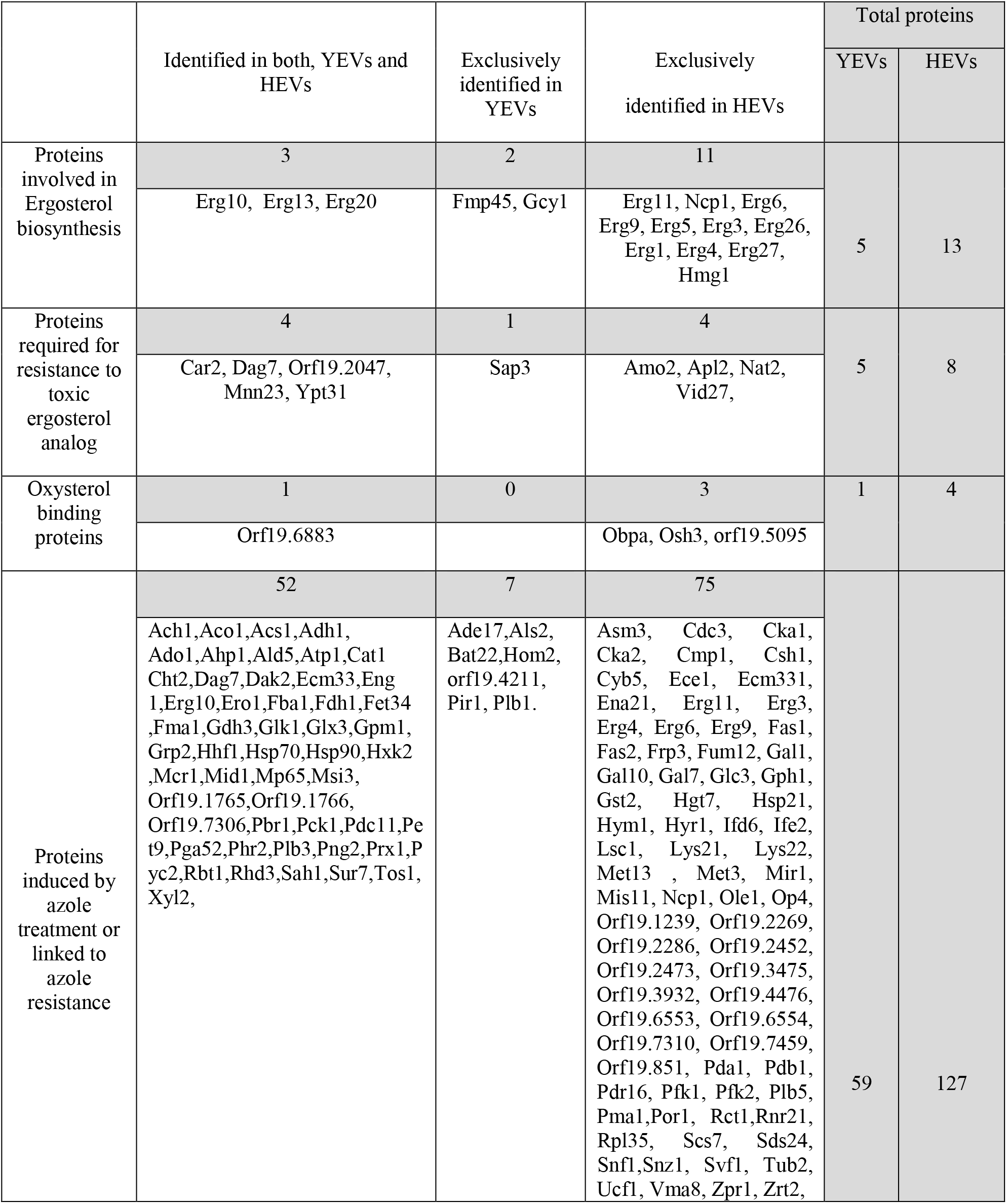
Proteins identified in YEVs and HEVs which are described in the CGD as involved in ergosterol biosynthesis, required for resistance to toxic ergosterol analog, induced by azole treatment or linked to azole resistance.

## Notes

### Competing Interest Statement

The authors have declared no competing interest.

## REFERENCES

1. Brown, G. D.; Denning, D. W.; Levitz, S. M., Tackling Human Fungal Infections. Science 2012, 336, (6082), 647–647.

2. Pappas, P. G.; Lionakis, M. S.; Arendrup, M. C.; Ostrosky-Zeichner, L.; Kullberg, B. J., Invasive candidiasis. Nat Rev Dis Primers 2018, 4, 18026.

3. Hofs, S.; Mogavero, S.; Hube, B., Interaction of Candida albicans with host cells: virulence factors, host defense, escape strategies, and the microbiota. J Microbiol 2016, 54, (3), 149–69.

4. Monteoliva, L.; Martinez-Lopez, R.; Pitarch, A.; Hernaez, M. L.; Serna, A.; Nombela, C.; Albar, J. P.; Gil, C., Quantitative proteome and acidic subproteome profiling of Candida albicans yeast-to-hypha transition. J Proteome Res 2011, 10, (2), 502–17.

5. Martinez-Gomariz, M.; Perumal, P.; Mekala, S.; Nombela, C.; Chaffin, W. L.; Gil, C., Proteomic analysis of cytoplasmic and surface proteins from yeast cells, hyphae, and biofilms of Candida albicans. Proteomics 2009, 9, (8), 2230–2252.

6. Hernaez, M. L.; Ximenez-Embun, P.; Martinez-Gomariz, M.; Gutierrez-Blazquez, M. D.; Nombela, C.; Gil, C., Identification of Candida albicans exposed surface proteins in vivo by a rapid proteomic approach. J Proteomics 2010, 73, (7), 1404–9.

7. Gil-Bona, A.; Parra-Giraldo, C. M.; Hernaez, M. L.; Reales-Calderon, J. A.; Solis, N. V.; Filler, S. G.; Monteoliva, L.; Gil, C., Candida albicans cell shaving uncovers new proteins involved in cell wall integrity, yeast to hypha transition, stress response and host-pathogen interaction. Journal of Proteomics 2015, 127, 340–351.

8. Marin, E.; Parra-Giraldo, C. M.; Hernandez-Haro, C.; Hernaez, M. L.; Nombela, C.; Monteoliva, L.; Gil, C., Candida albicans Shaving to Profile Human Serum Proteins on Hyphal Surface. Front Microbiol 2015, 6, 1343.

9. Gil-Bona, A.; Monteoliva, L.; Gil, C., Global Proteomic Profiling of the Secretome of Candida albicans ecm33 Cell Wall Mutant Reveals the Involvement of Ecm33 in Sap2 Secretion. Journal of Proteome Research 2015, 14, (10), 4270–4281.

10. Gil-Bona, A.; Amador-Garcia, A.; Gil, C.; Monteoliva, L., The external face of Candida albicans: A proteomic view of the cell surface and the extracellular environment. Journal of Proteomics 2018, 180, 70–79.

11. Chaffin, W. L.; Lopez-Ribot, J. L.; Casanova, M.; Gozalbo, D.; Martinez, J. P., Cell wall and secreted proteins of Candida albicans: identification, function, and expression. Microbiol Mol Biol Rev 1998, 62, (1), 130–80.

12. Lopez-Villar, E.; Monteoliva, L.; Larsen, M. R.; Sachon, E.; Shabaz, M.; Pardo, M.; Pla, J.; Gil, C.; Roepstorff, P.; Nombela, C., Genetic and proteomic evidences support the localization of yeast enolase in the cell surface. Proteomics 2006, 6 Suppl 1, S107–18.

13. Pitarch, A.; Sanchez, M.; Nombela, C.; Gil, C., Sequential fractionation and two-dimensional gel analysis unravels the complexity of the dimorphic fungus Candida albicans cell wall proteome. Mol Cell Proteomics 2002, 1, (12), 967–82.

14. Nombela, C.; Gil C Fau - Chaffin, W. L., Chaffin, W. L.; Non-conventional protein secretion in yeast. 2006, (0966–842X (Print)).

15. Rodrigues, M. L.; Nakayasu, E. S.; Almeida, I. C.; Nimrichter, L., The impact of proteomics on the understanding of functions and biogenesis of fungal extracellular vesicles. Journal of Proteomics 2014, 97, 177–186.

16. Anderson, J.; Mihalik, R.; Soll, D. R., Ultrastructure and antigenicity of the unique cell wall pimple of the Candida opaque phenotype. J Bacteriol 1990, 172, (1), 224–35.

17. Rodrigues, M. L.; Nimrichter L Fau - Oliveira, D. L., Oliveira Dl Fau - Frases, S., Frases S Fau - Miranda, K., Miranda K Fau - Zaragoza, O., Zaragoza O Fau - Alvarez, M., Alvarez M Fau - Nakouzi, A., Nakouzi A Fau - Feldmesser, M., Feldmesser M Fau - Casadevall, A., Casadevall, A.; Vesicular polysaccharide export in Cryptococcus neoformans is a eukaryotic solution to the problem of fungal trans-cell wall transport. 2007, (1535–9778 (Print)).

18. Albuquerque, P. C.; Nakayasu, E. S.; Rodrigues, M. L.; Frases, S.; Casadevall, A.; Zancope-Oliveira, R. M.; Almeida, I. C.; Nosanchuk, J. D., Vesicular transport in Histoplasma capsulatum: an effective mechanism for trans-cell wall transfer of proteins and lipids in ascomycetes. Cellular Microbiology 2008, 10, (8), 1695–1710.

19. Gil-Bona, A.; Llama-Palacios, A.; Parra, C. M.; Vivanco, F.; Nombela, C.; Monteoliva, L.; Gil, C., Proteomics Unravels Extracellular Vesicles as Carriers of Classical Cytoplasmic Proteins in Candida albicans. Journal of Proteome Research 2015, 14, (1), 142–153.

20. Vargas, G.; Rocha, J. D. B.; Oliveira, D. L.; Albuquerque, P. C.; Frases, S.; Santos, S. S.; Nosanchuk, J. D.; Gomes, A. M. O.; Medeiros, L. C. A. S.; Miranda, K.; Sobreira, T. J. P.; Nakayasu, E. S.; Arigi, E. A.; Casadevall, A.; Guimaraes, A. J.; Rodrigues, M. L.; Freire-de-Lima, C. G.; Almeida, I. C.; Nimrichter, L., Compositional and immunobiological analyses of extracellular vesicles released by Candida albicans. Cellular Microbiology 2015, 17, (3), 389–407.

21. Wolf, J. M.; Espadas, J.; Luque-Garcia, J.; Reynolds, T.; Casadevall, A., Lipid Biosynthetic Genes Affect Candida albicans Extracellular Vesicle Morphology, Cargo, and Immunostimulatory Properties. Eukaryotic Cell 2015, 14, (8), 745–754.

22. Malloci, M.; Perdomo, L.; Veerasamy, M.; Andriantsitohaina, R.; Simard, G.; Martinez, M. C., Extracellular Vesicles: Mechanisms in Human Health and Disease. Antioxidants & Redox Signaling 2019, 30, (6), 813–856.

23. Maas, S. L. N.; Breakefield, X. O.; Weaver, A. M., Extracellular Vesicles: Unique Intercellular Delivery Vehicles. Trends in Cell Biology 2017, 27, (3), 172–188.

24. Raposo, G.; Stoorvogel, W., Extracellular vesicles: Exosomes, microvesicles, and friends. Journal of Cell Biology 2013, 200, (4), 373–383.

25. Rodrigues, M. L.; Nakayasu, E. S.; Oliveira, D. L.; Nimrichter, L.; Nosanchuk, J. D.; Almeida, I. C.; Casadevall, A., Extracellular vesicles produced by Cryptococcus neoformans contain protein components associated with virulence. Eukaryotic Cell 2008, 7, (1), 58–67.

26. Zarnowski, R.; Sanchez, H.; Covelli, A. S.; Dominguez, E.; Jaromin, A.; Berhardt, J.; Heiss, C.; Azadi, P.; Mitchell, A.; Andes, D. R. A.-O. h. o. o.; Candida albicans biofilm-induced vesicles confer drug resistance through matrix biogenesis. 2018, (1545–7885 (Electronic)).

27. Zhao, K. N.; Bleackley, M.; Chisanga, D.; Gangoda, L.; Fonseka, P.; Liem, M.; Kalra, H.; Al Saffar, H.; Keerthikumar, S.; Ang, C. S.; Adda, C. G.; Jiang, L. Z.; Yap, K.; Poon, I. K.; Lock, P.; Bulone, V.; Anderson, M.; Mathivanan, S., Extracellular vesicles secreted by Saccharomyces cerevisiae are involved in cell wall remodelling. Communications Biology 2019, 2.

28. Freitas, M. S.; Bonato, V. L. D.; Pessoni, A. M.; Rodrigues, M. L.; Casadevall, A.; Almeidaa, F., Fungal Extracellular Vesicles as Potential Targets for Immune Interventions. Msphere 2019, 4, (6).

29. Herkert, P. F.; Amatuzzi, R. F.; Alves, L. R.; Rodrigues, M. L.; Extracellular Vesicles as Vehicles for the Delivery of Biologically Active Fungal Molecules. 2019, (1875–5550 (Electronic)).

30. Gillum, A. M.; Tsay, E. Y.; Kirsch, D. R., Isolation of the Candida albicans gene for orotidine-5’-phosphate decarboxylase by complementation of S. cerevisiae ura3 and E. coli pyrF mutations. Mol Gen Genet 1984, 198, (2), 179–82.

31. Sorgo, A. G.; Heilmann, C. J.; Dekker, H. L.; Brul, S.; de Koster, C. G.; Klis, F. M., Mass spectrometric analysis of the secretome of Candida albicans. Yeast 2010, 27, (8), 661–72.

32. Luo, T.; Kruger, T.; Knupfer, U.; Kasper, L.; Wielsch, N.; Hube, B.; Kortgen, A.; Bauer, M.; Giamarellos-Bourboulis, E. J.; Dimopoulos, G.; Brakhage, A. A.; Kniemeyer, O., Immunoproteomic Analysis of Antibody Responses to Extracellular Proteins of Candida albicans Revealing the Importance of Glycosylation for Antigen Recognition. Journal of Proteome Research 2016, 15, (8), 2394–2406.

33. Zybailov, B. L.; Florens, L.; Washburn, M. P., Quantitative shotgun proteomics using a protease with broad specificity and normalized spectral abundance factors. Molecular Biosystems 2007, 3, (5), 354–360.

34. Pathan, M.; Keerthikumar, S.; Chisanga, D.; Alessandro, R.; Ang, C. S.; Askenase, P.; Batagov, A. O.; Benito-Martin, A.; Camussi, G.; Clayton, A.; Collino, F.; Di Vizio, D.; Falcon-Perez, J.; Fonseca, P.; Fonseka, P.; Fontana, S.; Gho, Y. S.; Hendrix, A.; Nolte-’T Hoen, E.; Iraci, N.; Kastaniegaard, K.; Kislinger, T.; Kowal, J.; Kurochkin, I. V.; Leonardi, T.; Liang, Y.; Llorente, A.; Lunavat, T. R.; Maji, S.; Monteleone, F.; Overbye, A.; Panaretakis, T.; Patel, T.; Peinado, H.; Pluchino, S.; Principe, S.; Ronquist, G.; Royo, F.; Sahoo, S.; Spinelli, C.; Stensballe, A.; Thery, C.; van Herwijnen, M. J. C.; Wauben, M.; Welton, J. L.; Zhao, K.; Mathivanan, S., A novel community driven software for functional enrichment analysis of extracellular vesicles data. Journal of Extracellular Vesicles 2017, 6.

35. Kanehisa, M.; Goto, S., KEGG: Kyoto Encyclopedia of Genes and Genomes. Nucleic Acids Research 2000, 28, (1), 27–30.

36. Xu, Y.; Sheng, F.; Zhao, J.; Chen, L.; Li, C.; ERG11 mutations and expression of resistance genes in fluconazole-resistant Candida albicans isolates. 2015, (1432–072X (Electronic)).

37. Xu, D. M.; Jiang, B.; Ketela, T.; Lemieux, S.; Veillette, K.; Martel, N.; Davison, J.; Sillaots, S.; Trosok, S.; Bachewich, C.; Bussey, H.; Youngman, P.; Roemer, T., Genome-wide fitness test and mechanism-of-action studies of inhibitory compounds in Candida albicans. Plos Pathogens 2007, 3, (6), 835–848.

38. Huang, S. H.; Wu, C. H.; Chang, Y. C.; Kwon-Chung, K. J.; Brown, R. J.; Jong, A., Cryptococcus neoformans-Derived Microvesicles Enhance the Pathogenesis of Fungal Brain Infection. Plos One 2012, 7, (11).

39. Dawson, C. S.; Garcia-Ceron, D.; Rajapaksha, H.; Faou, P.; Bleackley, M. R.; Anderson, M. A., Protein markers for Candida albicans EVs include claudin-like Sur7 family proteins. Journal of Extracellular Vesicles 2020, 9, (1).

40. Moyes, D. L.; Wilson, D.; Richardson, J. P.; Mogavero, S.; Tang, S. X.; Wernecke, J.; Höfs, S.; Gratacap, R. L.; Robbins, J.; Runglall, M.; Murciano, C.; Blagojevic, M.; Thavaraj, S.; Förster, T. M.; Hebecker, B.; Kasper, L.; Vizcay, G.; Iancu, S. I.; Kichik, N.; Häder, A.; Kurzai, O.; Luo, T.; Krüger, T.; Kniemeyer, O.; Cota, E.; Bader, O.; Wheeler, R. T.; Gutsmann, T.; Hube, B.; Naglik, J. R., Candidalysin is a fungal peptide toxin critical for mucosal infection. Nature 2016, 532, (7597), 64–68.

41. Naglik, J. R.; Gaffen, S. L.; Hube, B., Candidalysin: discovery and function in Candida albicans infections. Current Opinion in Microbiology 2019, 52, 100–109.

42. Bader, O.; Krauke, Y.; Hube, B., Processing of predicted substrates of fungal Kex2 proteinases from Candida albicans, C. glabrata, Saccharomyces cerevisiae and Pichia pastoris. BMC Microbiology 2008, 8, (1), 116.

43. Richardson, J. P.; Mogavero, S.; Moyes, D. L.; Blagojevic, M.; Krüger, T.; Verma, A. H.; Coleman, B. M.; De La Cruz Diaz, J., Schulz, D.; Ponde, N. O.; Carrano, G.; Kniemeyer, O.; Wilson, D.; Bader, O.; Enoiu, S. I.; Ho, J.; Kichik, N.; Gaffen, S. L.; Hube, B.; Naglik, J. R., Processing of Candida albicans Ece1p Is Critical for Candidalysin Maturation and Fungal Virulence. mBio 2018, 9, (1).

44. Vialas, V.; Perumal, P.; Gutierrez, D.; Ximenez-Embun, P.; Nombela, C.; Gil, C.; Chaffin, W. L., Cell surface shaving of Candida albicans biofilms, hyphae, and yeast form cells. Proteomics 2012, 12, (14), 2331–2339.

45. Vialas, V.; Sun, Z.; Reales-Calderon, J. A.; Hernaez, M. L.; Casas, V.; Carrascal, M.; Abian, J.; Monteoliva, L.; Deutsch, E. W.; Moritz, R. L.; Gil, C., A comprehensive Candida albicans PeptideAtlas build enables deep proteome coverage. J Proteomics 2016, 131, 122–130.

46. Kasper, L.; Konig, A.; Koenig, P. A.; Gresnigt, M. S.; Westman, J.; Drummond, R. A.; Lionakis, M. S.; Gross, O.; Ruland, J.; Naglik, J. R.; Hube, B., The fungal peptide toxin Candidalysin activates the NLRP3 inflammasome and causes cytolysis in mononuclear phagocytes. Nature Communications 2018, 9.

47. Gropp, K.; Schild, L.; Hube, B.; Zipfel, P. F.; Skerka, C., Secreted aspartic proteinases (Saps) of Candida albicans degrade host complement. Molecular Immunology 2009, 46, (14), 2835–2836.

48. Felk, A.; Kretschmar M Fau - Albrecht, A., Albrecht A Fau - Schaller, M., Schaller M Fau - Beinhauer, S., Beinhauer S Fau - Nichterlein, T., Nichterlein T Fau - Sanglard, D., Sanglard D Fau - Korting, H. C., Korting Hc Fau - Schäfer, W.; Schäfer W Fau - Hube, B., Hube, B.; Candida albicans hyphal formation and the expression of the Efg1-regulated proteinases Sap4 to Sap6 are required for the invasion of parenchymal organs. 2002, (0019–9567 (Print)).

49. Albrecht, A.; Felk, A.; Pichova, I.; Naglik, J. R.; Schaller, M.; de Groot, P.; MacCallum, D.; Odds, F. C.; Schafer, W.; Klis, F.; Monod, M.; Hube, B., Glycosylphosphatidylinositol-anchored proteases of Candida albicans target proteins necessary for both cellular processes and host-pathogen interactions. Journal of Biological Chemistry 2006, 281, (2), 688–694.

50. Knechtle, P.; Goyard, S.; Brachat, S.; Ibrahim-Granet, O.; d’Enfert, C., Phosphatidylinositol-dependent phospholipases C Plc2 and Plc3 of Candida albicans are dispensable for morphogenesis and host-pathogen interaction. Res Microbiol 2005, 156, (7), 822–9.

51. Phan, Q. T.; Myers, C. L.; Fu, Y.; Sheppard, D. C.; Yeaman, M. R.; Welch, W. H.; Ibrahim, A. S.; Edwards, J. E.; Filler, S. G., Als3 is a Candida albicans invasin that binds to cadherins and induces endocytosis by host cells. Plos Biology 2007, 5, (3), 543–557.

52. Zhu, W. D.; Phan, Q. T.; Boontheung, P.; Solis, N. V.; Loo, J. A.; Filler, S. G., EGFR and HER2 receptor kinase signaling mediate epithelial cell invasion by Candida albicans during oropharyngeal infection. Proceedings of the National Academy of Sciences of the United States of America 2012, 109, (35), 14194–14199.

53. Moyes, D. L.; Richardson, J. P.; Naglik, J. R., Candida albicans-epithelial interactions and pathogenicity mechanisms: scratching the surface. Virulence 2015, 6, (4), 338–46.

54. Martinez-Lopez, R.; Nombela, C.; Diez-Orejas, R.; Monteoliva, L.; Gil, C., Immunoproteomic analysis of the protective response obtained from vaccination with Candida albicans ecm33 cell wall mutant in mice. Proteomics 2008, 8, (13), 2651–64.

55. Hershko, A.; Ciechanover, A., The ubiquitin system. Annual Review of Biochemistry 1998, 67, 425-479.

56. Nandi, D.; Tahiliani, P.; Kumar, A.; Chandu, D., The ubiquitin-proteasome system. Journal of Biosciences 2006, 31, (1), 137–155.

57. Hossain, S. A.-O.; Veri, A. A.-O.; Cowen, L. A.-O.; The Proteasome Governs Fungal Morphogenesis via Functional Connections with Hsp90 and cAMP-Protein Kinase A Signaling. LID - 10.1128/mBio.00290-20 [doi] LID - e00290-20. 2020, (2150–7511 (Electronic)).

58. Lin, W. C.; Tsai, C. Y.; Huang, J. M.; Wu, S. R.; Chu, L. J.; Huang, K. Y., Quantitative proteomic analysis and functional characterization of Acanthamoeba castellanii exosome-like vesicles. Parasites & Vectors 2019, 12, (1).

59. Jiang, L.; Zhao J Fau - Guo, R., Guo R Fau - Li, J., Li J Fau - Yu, L., Yu L Fau - Xu, D., Xu, D.; Functional characterization and virulence study of ADE8 and GUA1 genes involved in the de novo purine biosynthesis in Candida albicans. (1567–1364 (Electronic)).

60. Johnstone, R. M.; Adam, M.; Hammond, J. R.; Orr, L.; Turbide, C., Vesicle Formation during Reticulocyte Maturation - Association of Plasma-Membrane Activities with Released Vesicles (Exosomes). Journal of Biological Chemistry 1987, 262, (19), 9412–9420.

61. Kowal, J.; Arras, G.; Colombo, M.; Jouve, M.; Morath, J. P.; Primdal-Bengtson, B.; Dingli, F.; Loew, D.; Tkach, M.; Thery, C., Proteomic comparison defines novel markers to characterize heterogeneous populations of extracellular vesicle subtypes. Proceedings of the National Academy of Sciences of the United States of America 2016, 113, (8), E968–E977.

62. Colombo, M.; Raposo, G.; Thery, C., Biogenesis, Secretion, and Intercellular Interactions of Exosomes and Other Extracellular Vesicles. Annual Review of Cell and Developmental Biology, Vol 30 2014, 30, 255–289.

63. Oliveira, D. L.; Nakayasu, E. S.; Joffe, L. S.; Guimaraes, A. J.; Sobreira, T. J. P.; Nosanchuk, J. D.; Cordero, R. J. B.; Frases, S.; Casadevall, A.; Almeida, I. C.; Nimrichter, L.; Rodrigues, M. L., Characterization of Yeast Extracellular Vesicles: Evidence for the Participation of Different Pathways of Cellular Traffic in Vesicle Biogenesis. Plos One 2010, 5, (6).

